# CoNECo: A Corpus for Named Entity recognition and normalization of protein Complexes

**DOI:** 10.1101/2024.05.18.594800

**Authors:** Katerina Nastou, Mikaela Koutrouli, Sampo Pyysalo, Lars Juhl Jensen

**Affiliations:** Novo Nordisk Foundation Center for Protein Research, University of Copenhagen, Blegdamsvej 3, 2200, Copenhagen, Denmark; TurkuNLP Group, Department of Computing, University of Turku, Turku, Finland

**Keywords:** Named Entity Recognition, Transformer-based models, Dictionary-based NER, Protein-containing complex

## Abstract

**Motivation:** Despite significant progress in biomedical information extraction, there is a lack of resources for Named Entity Recognition (NER) and Normalization (NEN) of protein-containing complexes. Current resources inadequately address the recognition of protein-containing complex names across different organisms, underscoring the crucial need for a dedicated corpus.

**Results:** We introduce the Complex Named Entity Corpus (CoNECo), an annotated corpus for NER and NEN of complexes. CoNECo comprises 1,621 documents with 2,052 entities, 1,976 of which are normalized to Gene Ontology. We divided the corpus into training, development, and test sets and trained both a transformer-based and dictionary-based tagger on them. Evaluation on the test set demonstrated robust performance, with F1-scores of 73.7% and 61.2%, respectively. Subsequently, we applied the best taggers for comprehensive tagging of the entire openly accessible biomedical literature.

**Availability:** All resources, including the annotated corpus, training data, and code, are available to the community through Zenodo https://zenodo.org/records/11263147 and GitHub https://zenodo.org/records/10693653.

## Introduction

Improved deep-learning methodologies [Milošević and Thielemann, 2023], such as Transformer-based models [Vaswani et al., 2017] pre-trained on large corpora, coupled with efforts towards annotation of biomedical text corpora [Luoma et al., 2023, Luo et al., 2022, Li et al., 2016, Herrero-Zazo et al., 2013, Krallinger et al., 2008, Pyysalo et al., 2007, Kim et al., 2003], have recently led to major advances in the field of information extraction. The aforementioned corpora have facilitated the development of methods that can accurately recognize a variety of entity types, including chemicals [Krallinger et al., 2015], genes/proteins [Smith et al., 2008], organisms [Luoma et al., 2023], and diseases [Doğan et al., 2014]. Leveraging these resources, deep learning-based methods have made great progress in named entity recognition tasks for these entities [Lee et al., 2020].

Despite their biological and pharmacological importance [Santos et al., 2017, Harding et al., 2024], there remains a conspicuous lack of a corpus specifically designed to evaluate Named Entity Recognition (NER) and Normalization (NEN) of protein-containing complexes (henceforth referred to as `complex`). While there are resources available for recognition of human `complex` names [Bachman et al., 2018], or annotation of mentions of type `complex` as part of a BioNLP Shared Task focused on Relation Extraction [Bossy et al., 2015], the development of a comprehensive corpus that facilitates the training and evaluation of dictionary-based or deep learning-based NER systems for complex names across multiple organisms is notably absent. This gap highlights a critical area of need, given the biological importance of complexes.

In this work we propose the annotation of a new corpus, named CoNECo (Complex Named Entity Corpus), to serve the purposes of NER and NEN of protein complexes. Specifically, we have annotated 1,621 documents with 2,052 `complex` named entities, normalized in Gene Ontology (GO) [Ashburner et al., 2000, Aleksander et al., 2023]. We have split the CoNECo documents into training, development, and test sets. We used the training and development sets to train a transformer-based tagger [Luoma et al., 2023] and to improve a dictionary-based tagger [Pafilis et al., 2013]. We evaluated both taggers on the held-out test set, achieving F1-scores of 73.7% and 61.2%, respectively, and used the best taggers for large-scale tagging of all PubMed abstracts and open-access articles from PubMed Central. All data used and produced in this project, along with the code to reproduce the results, are openly accessible via Zenodo and GitHub.

## Materials and methods

### The CoNECo corpus

#### Document selection

The first step towards the generation of a corpus for the annotation of `complex` named entities was document selection. To benefit from existing resources we initially focused our efforts on previously annotated corpora where work was already done for the annotation of `complex`, which largely fitted the definition introduced above. As complexes play crucial roles in cellular signaling, we decided to improve the balance of the corpus by expanding it with documents related to this topic.

The selection of the documents consisted of three steps which are detailed below:

1. ComplexTome corpus [Mehryary et al., 2023]: All documents from ComplexTome, a corpus designed for training a deep learning-based relation extraction system for physical molecular interactions were selected. These documents contained named entity annotations for complexes, fitting with our definition, and were kept during annotation. Moreover, since this is a corpus for extraction of physical protein interactions, it inherently pertains to the topic of protein complexes.
2. Expansion with 100 additional Reactome abstracts: As the ComplexTome contains 300 documents used for the annotation of pathways in the Reactome pathway knowledgebase [Gillespie et al., 2022], we selected 100 extra abstracts from Reactome to increase the representation of signaling-related documents in CoNECo.
3. Event Extraction for Post-Translational Modifications corpus from the BioNLP 2010 workshop [Ohta et al., 2010]: 234 signaling-related abstracts were selected from the 388 total abstracts in this corpus based on the existence of at least one post-translational modification event and more than one entity in a document.

The corpus was split into training, development, and test sets, keeping the original split for ComplexTome and assigning the extra 334 documents to keep a 60%/20%/20% document-level split. All documents in CoNECo were annotated within the BRAT rapid annotation tool [Stenetorp et al., 2012].

#### Named entity annotation

CoNECo is a corpus aiming at NER and NEN of protein-containing complexes. As such, there is a single annotated entity type, namely “protein-containing complex”. To annotate and normalize such entities in text, we primarily built on GO [Ashburner et al., 2000, Aleksander et al., 2023]. GO has a substantial representation of protein-containing complexes, rooted at the sub-ontology of the homonymous GO term (*GO:0032991*) in the GO Cellular Component (GO:CC) ontology, and contains 2103 terms. The definition of these entities can be summarized as *a stable set of interacting proteins which can be co-purified by an acceptable method, and where the complex has been shown to exist as an isolated, functional unit in vivo*. An example annotation is given in Figure 1.

**Fig. 1.**
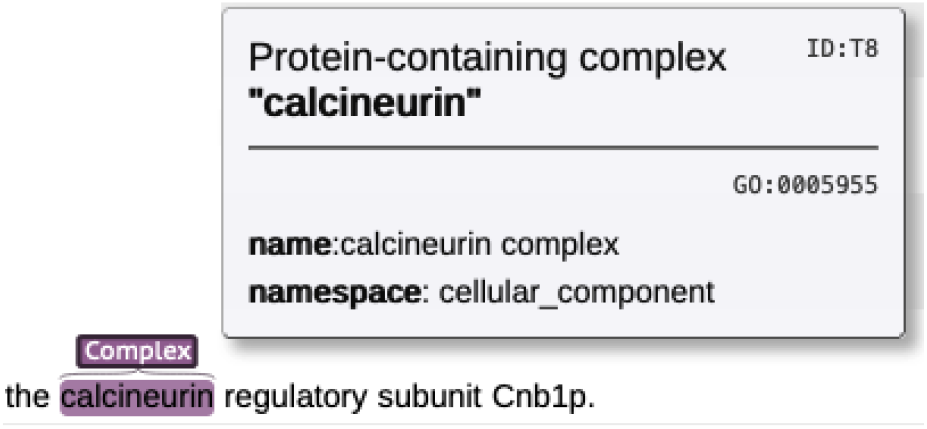
Illustration of a complex entity representation in CoNECo. The named entity (“calcineurin”) has been identified by the annotators and normalized to the *calcineurin complex* term in GO (*GO:0005955*).

It should be noted that NEN is decoupled from NER, allowing for the annotation of entity mentions that align with the definition introduced above, even if they are not present in GO, and thus cannot be normalized. This approach enriches the corpus with a more comprehensive annotation, enhancing its usability across various applications. Our decision to normalize to GO, as opposed to other specialized resources like Complex Portal [Meldal et al., 2022], is driven by the widespread adoption and organism-agnostic nature of GO, making it a versatile and universally applicable standard in the community. Our previous experience in annotating a deep learning-ready corpus for NER [Luoma et al., 2023], has highlighted the importance of boundary consistency in entity annotations. Extra care has also been taken in CoNECo, to ensure that the minimal span containing the full name of the entity mentioned in the text is annotated and that the marked span starts and ends on a boundary between an alphanumeric string and a non-alphanumeric character. When a `complex` name coincides with the name of a protein or its protein constituents or the name of a protein family, separated by any punctuation, the `complex` entity is not annotated. Moreover, no overlapping annotations between `complex` names are allowed, and the leftmost longest matching entity is annotated in cases where there are overlaps between different `complex` names.

Another important factor to ensure the quality of the corpus is ensuring the consistency of mentions in text and the names and synonyms that these normalize to. We have performed a semi-automated check to evaluate this consistency, which produced a list of `complex` names that do not have a clear match in GO. All these names were manually checked against alternative Complex Portal [Meldal et al., 2022] and Reactome [Gillespie et al., 2022], to assess whether a link to GO entries could be obtained via them. If this was still not possible, but a link to alternative resources or the literature could be found, a comment was left in the corpus, and the entities were annotated solely for NER.

To evaluate the general quality of annotations, we randomly selected approximately 5% of abstracts from the training set and provided them to two curators in two rounds for independent annotation. Subsequently, we calculated the F-score of their agreement to determine the consistency of annotations and the overall corpus quality.

Detailed annotation guidelines are provided through Zenodo and this link https://katnastou.github.io/annodoc-CoNECo/.

In addition to human annotators we have examined the possibility of using an annotation assistant for annotating documents for the CoNECo corpus by providing it with the annotation guidelines. To assess if this is possible we decided to use the documents from the second round of inter-annotator agreement (IAA) and provide those as prompts to custom versions of ChatGPT 4.0. The set of prompts is available in Supplementary Section 1.

### Dictionary-based NER and NEN

The JensenLab tagger [Jensen, 2016] is a fast, dictionary-based method for the recognition and normalization of several biomedical entity types. Since dictionary-based tagging is widely used and adopted in several biological resources — including for tagging protein and organism names for the influential database of protein–protein interactions STRING [Szklarczyk et al., 2023] — we wanted to ensure that the CoNECo corpus is suitable for evaluating dictionary-based methods, like the JensenLab tagger. We built a dictionary for tagger using all terms of GO:CC below “protein-containing complex”. Additional `complex` names from Complex Portal were added to the dictionary following the process below:

1. The mapping between entries in Complex Portal and `complex` terms in Gene Ontology was obtained by properly filtering the mapping file available through the Complex Portal FTP http://ftp.ebi.ac.uk/pub/databases/intact/complex/current/go/complex_portal.v2.gpad
2. Complex Portal entries with multiple mappings to GO entries were ignored, after manual checking of a random sample of these entries, where several cases where synonyms seemed to match only one of the mapped entries were identified.
3. Complex Portal names that were already present in GO were removed.
4. Complex Portal names that included names of their protein constituents were removed either automatically or manually and the final list of extra names was obtained.

The dictionary was expanded with orthographic variants of the names coming from GO:CC and those added from Complex Portal. Specifically, forms where the word “complex” is not included were generated as well as plural and adjectival forms of existing names. Orthographic variation related to hyphenation or spacing is handled internally by the JensenLab tagger as described in Pafilis et al. [2013] and no such variants were thus added.

The JensenLab tagger software was run on the combined training and development set to identify potential issues with the dictionary generated following the process above and the dictionary files — including the blocklist — were updated accordingly.

After the dictionary build was complete, the dictionary-based tagger was run on the corpus test set and an evaluation of both NER and NEN was performed.

### Transformer-based NER

Despite the widespread use of dictionary-based methods, it would be an omission not to assess whether the corpus is also useful for deep learning-based NER, considering that the majority of the biomedical text-mining community has now migrated to such methods, especially Transformer-based ones [Miranda-Escalada et al., 2023]. We have selected the RoBERTa-large-PM-M3-Voc (hereafter RoBERTa-biolm) [Lewis et al., 2020] model, as it has been repeatedly shown to outperform all other models in biomedical tasks [Miranda-Escalada et al., 2023, Luoma et al., 2023].

We used the method described in [Luoma and Pyysalo, 2020] for training and evaluation of a RoBERTa-biolm model. We attached a single fully connected layer on top of the Transformer architecture and fine-tuned the model to detect `complex` entities in the training data by classifying individual tokens in input samples. We selected hyperparameters by doing a grid search with three repetitions for each parameter set — to minimize the effect of initial random weights on evaluation scores [Mehryary et al., 2016] — training on the documents in the training set, and validating on those in the development set. The best mean F1-score on the development set was used to select the set of hyperparameters for training the model for evaluation on the test set. All training and development data are used in fine-tuning the model with the best set of hyperparameters. The process was repeated three times and the results shown in this paper are expressed as a mean and standard deviation of the exact and overlapping match F1-score. The latter allows us to compare the performance with the dictionary-based tagger.

## Results and discussion

### Corpus statistics

CoNECo comprises 1,621 documents with a total of 398,718 words (calculated using the BERT pre-tokenizer) and a total number of 2,052 `complex` named entities 443 of which are unique names. From a first glance at these numbers, the density of annotated mentions in CoNECo (0.5%) is low in comparison to other biomedical NER corpora. Specifically, we compared the density to S1000 [Luoma et al., 2023] which was 2.4% (6,328 total mentions in 262,293 words), BC2GM [Smith et al., 2008] with 4.3% (24,596 mentions in 569,912 words), BC5CDR [Li et al., 2016] with 4.4% and 3.5% density for drugs/chemicals and diseases, respectively (15,915 drug/chemical and 12,617 disease mentions in 360,373 words), and NCBI Disease [Doğan et al., 2014] with 3.7% density (6,892 mentions in 184,552 words). In all cases, the number of words in the corpus is calculated using the BERT pre-tokenizer. This level of sparsity in CoNECo means that precision could be severely affected for any tagging method, because mentions are rare, and thus result in lower F-scores for protein-containing `complex` recognition in comparison to other biomedical NER tasks. There is an additional challenge for machine learning-based methods, as the number of positive examples in the training set might not be sufficient to learn the real-world data distribution, and this could significantly affect performance on the test data.

Table 1 shows detailed statistics for the training, development, and test sets, as well as the entire corpus.

**Table 1.** CoNECo corpus statistics. The number of unique names and normalizations in the train, devel, and test sets does not sum up to the total number of unique names and normalizations in the corpus, as there can be different unique names and normalizations in each set, which are duplicates once the entire corpus is considered.

| Category | Total Documents | Total Mentions | Unique Names | Total Normalized Mentions | Unique Normalizations |
| --- | --- | --- | --- | --- | --- |
| Train set | 983 | 1360 | 330 | 1310 | 138 |
| Devel set | 320 | 409 | 126 | 387 | 63 |
| Test set | 318 | 283 | 103 | 279 | 58 |
| Total | 1621 | 2052 | 443 | 1976 | 183 |

We attained a 92.5% IAA after two rounds of IAA on 50 documents, showcasing the high quality of CoNECo. Additional guidelines were added between the first and second rounds, which allowed us to reach an above 90% IAA. After checking the differences in the second round all inconsistencies were attributed to annotation errors, which were fixed, and there was no need for a further update of the annotation guidelines. Moreover, the annotators were in full agreement about the normalization of entities to GO:CC from the first round of IAA.

### Using a custom ChatGPT as an annotator

To assess the possibility of automating document annotation for the task of `complex` NER, we have created 3 custom versions of ChatGPT. In the first version, we have provided the full annotation guidelines (with examples) as the instructions in the configuration of ChatGPT and created “CoNECo GPT - full” available through the Explore GPTs section of ChatGPT4.0 https://chat.openai.com/g/g-1uV7nfJTA-coneco-gpt-full. The instructions (Supplementary Section 2) correspond to the exact annotation guidelines provided to human annotators, with slight changes necessary to describe what was shown in markdown in the original documentation. We then provided the 25 prompts (Supplementary Section 1) one by one and within the same chat session. The terms identified as `complex` and how many of them are True Positives (TP), False Positives (FP), and False Negatives (FN) are shown in Supplementary Table 1. We used the human annotations in this set of 25 documents as a gold standard to calculate performance metrics, applying the overlapping matching criterion. The F-score for IAA with the human annotators in this experiment is 20.5% (Precision=12.5%, Recall=58.2%). The low F-score is mainly a result of the detection of several False Positives. Comparing that to the over 90% IAA between the human annotators it is evident that this custom version of ChatGPT cannot be used for annotation purposes.

To explore if providing the full annotation guidelines is what creates confusion to this model, we decided to provide a smaller set of instructions without examples. We have created “CoNECo GPT - small”, available through https://chat.openai.com/g/g-Ns0dcCn8c-coneco-gpt-small providing the set of instructions shown in Supplementary Section 3. The prompts remain the same and detailed results of using this version are provided in Supplementary Table 2. The F-score for IAA with the human annotators in this case is 30.5% (Precision=19.1%, Recall=75.5%). The results are better this time, but still much lower than the human annotator performance, to allow this to be a viable alternative.

**Table 2.** Error analysis for the JensenLab and the Transformer-based taggers.

| Error categories | Total |  | FN |  | FP |  |
| --- | --- | --- | --- | --- | --- | --- |
|  | dict-based | TF-based | dict-based | TF-based | dict-based | TF-based |
| Annotation error | 9 | 26 | 0 | 0 | 9 | 26 |
| Ambiguous name | 100 | 124 | 0 | 35 | 100 | 89 |
| Dictionary error | 112 | – | 97 | – | 15 | – |
| Discontinuous entity | 1 | 0 | 1 | 0 | 0 | 0 |
| Part-of longer entity | 13 | 4 | 0 | 0 | 13 | 4 |
| Unidentified model error | – | 13 | – | 13 | – | 0 |
| Total | 235 | 167 | 98 | 48 | 137 | 119 |

\*FN: false negative, FP: false positive
dict-based: dictionary-based tagger
TF-based:Transformer-based tagger

Since using fewer instructions during the creation of a custom ChatGPT provided slightly better results, we decided to perform one last experiment by creating a minimal set of instructions based on our annotation guidelines. In this case, we asked the non-customized version of ChatGPT 4.0 (https://chat.openai.com) to summarize the instructions for us. We have again provided the minimal set of instructions (Supplementary Section 4) to a custom ChatGPT, thus creating “CoNECo GPT - minimal” available via https://chat.openai.com/g/g-C6Nx12aEL-coneco-gpt-minimal. The full results for the same set of 25 prompts are shown in Supplementary Table 3. This minimal set of guidelines gives a significant boost to the IAA F-score, which now reaches 63.7%. This is mainly the result of a (*∼*60%) increase in Precision (80.6%), while the drop in Recall is around 20% (52.7%). Most false positives are cases where “CoNECo GPT - minimal” has not followed the guideline of “*If entities are separated by a dash, they are examined and reported separately* “and has produced results that could easily be filtered out to increase the precision even further. Nevertheless, multiple entities are still not detected, with the recall being (*∼*50%).

Additional experiments might be conducted to improve the recall of a custom ChatGPT, and further prompts could be provided to encourage it to identify the annotated entities. However, this falls outside the scope of evaluating ChatGPT’s potential as a substitute for human annotators in either annotating or expanding the corpus. Investing more effort in refining prompts or developing more sophisticated models is arguably not going to achieve the objective of lowering the costs associated with corpus development. One more observation that is relevant for assessing the use of such a model for corpora annotation is that in contrast to human annotators, the fewer guidelines we provide to ChatGPT the better it performs. This makes it prohibitive to use a similar approach for document annotation for even more difficult tasks such as relation extraction, where it will not be possible to limit the guidelines to the degree we have done here. Finally, It should be noted that starting a new chat and providing the same set of prompts could produce different results than those presented above.

### Dictionary-based NER

The original dictionary was built using all terms of GO:CC below “protein-containing complex” (GO:0032991) and 989 extra names were added from Complex Portal. We used this dictionary to run the JensenLab tagger on the combined training and development sets and quickly identified several issues with GO:CC terms that clash with our annotation guidelines. Specifically, 78 GO:CC terms were excluded based on manual review, as they do not represent individual complexes but rather groups of complexes with a common function (e.g. *GO:1902494, catalytic complex*) or localization (e.g. *GO:0140513 nuclear protein-containing complex*). The full list of excluded GO:CC terms is provided in the annotation guidelines documentation (https://katnastou.github.io/annodoc-CoNECo). On top of these, more specific issues were identified with clashes between `complex` names and protein names or `complex` names and protein family names, that required an update to our list of filtered names. After all the dictionary cleaning, we ran the tagger on the combined training and development set and obtained an F-score of 68.0% (precision 74.5%, recall 62.5%). All dictionary files to run the JensenLab tagger are available through Zenodo.

The dictionary was used to run the Jensenlab tagger on the CoNECo test set. Since the JensenLab tagger finds left-most longest matches of the names in its dictionary, the **overlapping matching** criterion is used to evaluate both this and the Transformer-based tagger. The tagger reached a precision of 57.5% (185/322), a recall of 65.4% (185/283) and an F-score of 61.2% on the test set. Our hypothesis that the sparsity of mentions in this corpus could severely affect the precision of the tagger in the test set, turned out to be true, as there is a 17.5% drop in precision from the dictionary run on the combined training and development sets to the test set. A detailed analysis and comparison of the recognition errors produced by both methods is presented in the section “Error analysis” below.

In our evaluation of the normalization for JensenLab tagger, an impressive 94.6% F-score was achieved, since 175 out of 185 normalizations for matching spans were found to be identical, underscoring the efficacy of our method in accurately mapping entities to their corresponding standardized terms.

The 10 instances where mismatches occurred are documented in *Supplementary Table 4*. It appears that the majority of these mismatches (80%) can be attributed to the tagging of names that are either more general or more specific than the expected normalization. This outcome, while not perfectly aligned with the intended normalization, is still acceptable. Among the remaining cases, one instance involves a conflicting narrow synonym in Gene Ontology that is less preferred to the annotated normalization and an annotation error, where we neglected to assign a normalization to a specific entity.

### Transformer-based NER

We fine-tuned a pre-trained RoBERTa-biolm using the training set of the CoNECo corpus and used the development set to identify a set of hyperparameters where the mean average F1-score is the highest on the task of `complex` NER. We obtained best results with models trained for 9 epochs, with a learning rate of 2E-5, a batch size of 2, and a maximum sequence length of 128. The F-score is 73.1% (std=0.27%), the precision is 80.5% (std=1.18%), and the recall is 67.0% (std=0.91%). A model trained using this set of hyperparameters is available through Zenodo.

We then tested the performance of this model against the CoNECo test set and obtained an F-score of F 73.7%, a precision of 66.3% (234/353), and a recall of 83.0% (235/283). The performance of this model on the dev and test sets is similar, in terms of F-score, but we can see once more that the precision is affected in the test set run (14% drop). Moreover, as we inferred from the sparsity statistics, the results for this task are worse in comparison to other biomedical NER tasks, where Transformer-based methods tend to reach above 90% F-score [Lee et al., 2020, Miranda-Escalada et al., 2023, Luoma et al., 2023].

We made an additional comparison to check the overlap of matches between the two NER methods in the CoNECo test set. We found 186 overlapping matches, 167 matches unique to the Transformer-based tagger, and 136 matches unique to the JensenLab tagger. If we make a union of all the matches and evaluate against the CoNECo test set we get a precision of 57.3%, a recall of 87.0%, and an F-score of 69.0%. If on the other hand, we use only the consensus of the two methods we get the following results: precision=89.6%, recall=50.8%, and F-score=64.8%. Including all results leads to an increase in recall while taking the more strict approach and using only the consensus of the two methods leads to an almost 90% precision, but a huge drop in recall to *∼*50%. In summary, if one needs normalization, the consensus of the two methods will yield better results than the dictionary-based tagger alone. On the other hand, if normalization is not needed best results are obtained by using the Transformer-based method alone. In the next section, we look into the errors produced by both methods and further assess the strengths and shortcomings of each approach.

### Error analysis

We have looked at the errors produced by both methods in detail and grouped them in categories as presented in Table 2. *Supplementary Table 5* and *Supplementary Table 6*, provide a detailed overview of errors for the JensenLab tagger and Transformer-based tagger, respectively.

There are three error categories that affect both methods. “Annotation errors” result in all cases in False Positives and thus refer to cases where the annotators have missed an entity annotation, that is otherwise correctly recognized by either the dictionary- or Transformer-based tagger. Recalculation of the performance of the JensenLab tagger without these errors results in a 1.5% increase in F-score (62.7%) due to an increased precision of 60.3%. The result is even more prominent for the Transformer-based tagger, where fixing the identified annotation errors results in 77.6% F-score due to a 72.8% precision. Both approaches face issues with “ambiguous names” that can denote either a protein-containing complex, a gene/protein or a family or large groups of proteins. According to our annotation guidelines, protein and family named entities have priority over `complex` during annotation, and thus the ambiguous entities have remained unannotated. However, given the inherent ambiguity of biomedical entity names in such cases, it becomes evident how these can severely impact the effectiveness of either method. One special subcategory of ambiguity is “part-of longer entity” errors, where longer more specific entities are either not part of the dictionary or have failed to be recognized by the Transformer-based method.

The other error categories can only be attributed to one of the two methods. “Dictionary errors” are responsible for approximately half of the mistakes made by the dictionary-based tagger. This error category indicates that the effectiveness of dictionary-based approaches hinges on the quality of their source dictionary. The main impact of dictionary errors is on recall, primarily due to the names missing from the dictionary. It is important to acknowledge that this problem stems from the fact that normalization is an inherent part of dictionary-based NER, and these cases would also challenge the Transformer-based method if results were normalized, and normalization relied on Gene Ontology as a source. Although there are strategies to mitigate these problems, the absence of names from the normalization source is a severely limiting factor.

While “discontinuous entity” could affect either method, in CoNECo it has only affected the JensenLab tagger. In general “discontinuous entities” pose significant challenges for dictionary-based methods and are one of the issues that deep learning-based methods can resolve better. Finally, “Unidentified model error” is an error category that affects only the Transformer-based tagger and refers to errors for which we could not properly interpret what has led the model into making these mistakes.

### Large-scale tagging

Results on tagging of over 36 million PubMed abstracts (as of January 2024) and 6 million articles from the PMC open access subset (as of November 2023) for both the JensenLab tagger and the Transformer-based method are provided via Zenodo. Tagging with the JensenLab tagger yielded 26,657,127 `complex` matches, covering 42,120 unique surface forms. Due to the nature of JensenLab tagger, these matches all map back to specific GO:CC cellular component entries, as GO was used to generate the dictionary for tagging. Considering the impressive results we got for normalization on the CoNECo test set (F-score=94.6%), we expect that mapping to be accurate also in the large-scale tagging. Tagging with the Transformer-based model produced a total of 19,654,180 matches, out of which 105,242 names are unique.

These results suggest that many synonyms are missing from the reference resources (GO, Complex Portal) used to generate the dictionary of `complex` names, which leads to a lower total count compared to the deep learning-based method. To investigate this, we compared the overlap of matches between the two systems. The total number of common matches between the two is 8,544,366 — which leaves 18,112,761 `complex` matches found only by JensenLab tagger and 11,109,814 found only by the Transformer-based method.

The most frequent common matches are unambiguous `complex` names, as expected from the consensus between two completely different approaches. The matches produced only by the Transformer-based tagger confirm that GO and Complex Portal lack synonyms for known complexes. Additionally, several ambiguous names (e.g. SNARE), which correspond to both `complex` and protein/protein family names, are unique to the Transformer-based tagger large-scale results because they are intentionally blocked in the dictionary-based tagger. While the results unique to the JensenLab tagger in many cases are correct, they also include the types of false positives already described in the error analysis. These could be a good starting point for manual curation to improve the block list of the dictionary-based tagger. The files with the frequencies for all matches for both methods can be found on Zenodo.

## Conclusions

In this work, we present CoNECo, the first corpus specifically designed for training and evaluating NER methods for `complex` recognition. The entities in the corpus are normalized to Gene Ontology, allowing for the evaluation of NEN methods on top of NER. CoNECo consists of 1,621 documents and 2,052 named entity annotations (1,976 normalized to GO), 443 of which are unique. Despite the sparsity of the corpus, it has been shown that it can be used for training and evaluating Transformer-based language models (F-score=73.7%, 77.6% correcting for annotation errors), as well as dictionary-based methods (F-score=61.2%, 62.7% correcting for annotation errors). Error analysis results have shown that ambiguity in entity names is the main issue faced by both dictionary- and deep learning-based methods. Moreover, dictionary-based methods are severely affected by the omission of a plethora of synonyms in reference resources like GO and Complex Portal.

## Supporting information

Supplementary Section

## Acknowledgements

We thank the CSC – IT Center for Science for generous computational resources.

## Data availability

All data utilized in this project, the code to replicate the results and large-scale tagging results of the biomedical literature are available under open licenses from Zenodo https://zenodo.org/records/11263147 and GitHub https://zenodo.org/records/10693653.

