## Supplementary Section for "CoNECo: A Corpus for Named Entity recognition and normalization of protein Complexes"

### Supplementary Section 1: The set of prompts we provided to the custom ChatGPTs

In this work we have tested whether it is possible to use an annotation assistant for annotating documents for the CoNECo corpus, providing proper guidelines. In order to assess if this is possible we decided to use the second set of 25 documents that we used to calculate the interannotator agreement (IAA) between human annotators and provide those as prompts to custom versions of ChatGPT 4.0.

I want you to check the document below with ID 10359895 and I want your output to have only a comma separated list of terms that you consider protein complexes based on the rules. If you think something is a protein complex but consists of affixes (e.g. - or /) and markers, even when they are part of the same syntactic word with a Complex name, as long as there is a separating nonalphanumeric character, do not report them. And remember to check if the terms you want to report are proteins or families.: Binding of YY1 and Oct1 to a novel element that downregulates expression of IL-5 in human T cells. BACKGROUND: IL-5 controls development of eosinophilia and has been shown to be involved in the pathogenesis of allergic diseases. In both atopic and nonatopic asthma, elevated IL-5 has been detected in peripheral blood and the airways. IL-5 is produced mainly by activated T cells, and its expression is regulated at the transcriptional level. OBJECTIVE: This study focuses on the functional analysis of the human IL-5 (hIL-5) promoter and characterization of cis -regulatory elements and transcription factors involved in the suppression of IL-5 transcription in T cells. METHODS: Methods used in this study include DNase I footprint assays, electrophoretic mobility shift assays, and functional analysis by mammalian cell transfection involving deletion analysis and site-directed mutagenesis. RESULTS: We identified 5 protein binding regions (BRs) located within the proximal hIL-5 promoter. Functional analysis indicates that the BRs are involved in control of hIL-5 promoter activity. Two of these regions, BR3 and BR4 located at positions -102 to -73, have not previously been described as regulators of IL-5 expression in T cells. We show that the BR3 sequence contains a novel negative regulatory element located at positions -90 to -79 of the hIL-5 promoter, which binds Oct1, octamer-like, and YY1 nuclear factors. Substitution mutations, which abolished binding of these proteins to the BR3 sequence, significantly increased hIL-5 promoter activity in activated T cells. CONCLUSION: We suggest that Oct1, YY1, and octamer-like factors binding to the -90/-79 sequence within the proximal IL-5 promoter are involved in suppression of IL-5 transcription in T cells.

I want you to check the document below with ID 10660620 and I want your output to have only a comma separated list of terms that you consider protein complexes based on the rules. If you think something is a protein complex but consists of affixes (e.g. - or /) and markers, even when they are part of the same syntactic word with a Complex name, as long as there is a separating nonalphanumeric character, do not report them. And remember to check if the terms you want to report are proteins or families.: PILRalpha, a novel immunoreceptor tyrosine-based inhibitory motif-bearing protein, recruits SHP-1 upon tyrosine phosphorylation and is paired with the truncated counterpart PILRbeta. SHP-1-mediated dephosphorylation of protein tyrosine residues is central to the regulation of several cell signaling pathways, the specificity of which is dictated by the intrinsic affinity of SH2 domains for the flanking sequences of phosphotyrosine residues. By using a modified yeast two-hybrid system and SHP-1 as bait, we have cloned a human cDNA, PILRalpha, encoding a 303-amino acid immunoglobulin-like transmembrane receptor bearing two cytoplasmic tyrosines positioned within an immunoreceptor tyrosine-based inhibitory motif. Substrate trapping in combination with pervanadate treatment of 293T cells confirms that PILRalpha associates with SHP-1 in vivo upon tyrosine phosphorylation. Mutation of the tyrosine residues in PILRalpha indicates the pivotal role of the Tyr-269 residue in recruiting SHP-1. Surface plasmon resonance analysis further suggests that the association between PILRalpha-Tyr-269 and SHP-1 is mediated primarily via the amino-terminal SH2 domain of the latter. Polymerase chain reaction amplification of cDNA in combination with genomic sequence analysis revealed a second gene, PILRbeta, coding for a putative activating receptor as suggested by a truncated cytoplasmic tail and a charged lysine residue in its transmembrane region. The PILRalpha and PILRbeta genes are localized to chromosome 7 which is in contrast with the mapping of known members of the inhibitory receptor superfamily.

I want you to check the document below with ID 11331609 and I want your output to have only a comma separated list of terms that you consider protein complexes based on the rules. If you think something is a protein complex but consists of affixes (e.g. - or /) and markers, even when they are part of the same syntactic word with a Complex name, as long as there is a separating nonalphanumeric character, do not report them. And remember to check if the terms you want to report are proteins or families.: Sharp, an inducible cofactor that integrates nuclear receptor repression and activation. A yeast two-hybrid screen using the conserved carboxyl terminus of the nuclear receptor corepressor SMRT as a bait led to the isolation of a novel human gene termed SHARP (SMRT/HDAC1 Associated Repressor Protein). SHARP is a potent transcriptional repressor whose repression domain (RD) interacts directly with SMRT and at least five members of the NuRD complex including HDAC1 and HDAC2. In addition, SHARP binds to the steroid receptor RNA coactivator SRA via an intrinsic RNA binding domain and suppresses SRA-potentiated steroid receptor transcription activity. Accordingly, SHARP has the capacity to modulate both liganded and nonliganded nuclear receptors. Surprisingly, the expression of SHARP is itself steroid inducible, suggesting a simple feedback mechanism for attenuation of the hormonal response.

I want you to check the document below with ID 11359852 and I want your output to have only a comma separated list of terms that you consider protein complexes based on the rules. If you think something is a protein complex but consists of affixes (e.g. - or /) and markers, even when they are part of the same syntactic word with a Complex name, as long as there is a separating nonalphanumeric character, do not report them. And remember to check if the terms you want to report are proteins or families.: Thiopalmitoylation of myelin proteolipid protein epitopes enhances immunogenicity and encephalitogenicity. Proteolipid protein (PLP) is the most abundant protein of CNS myelin, and is posttranslationally acylated by covalent attachment of long chain fatty acids to cysteine residues via a thioester linkage. Two of the acylation sites are within epitopes of PLP that are encephalitogenic in SJL/J mice (PLP(104-117) and PLP(139-151)) and against which increased immune responses have been detected in some multiple sclerosis patients. It is known that attachment of certain types of lipid side chains to peptides can result in their enhanced immunogenicity. The aim of this study was to determine whether thioacylated PLP peptides, as occur in the native protein, are more immunogenic than their nonacylated counterparts, and whether thioacylation influences the development of autoreactivity and experimental autoimmune encephalomyelitis. The results show that in comparison with nonacylated peptides, thioacylated PLP lipopeptides can induce greater T cell and Ab responses to both the acylated and nonacylated peptides. They also enhanced the development and chronicity of experimental autoimmune encephalomyelitis. Synthetic peptides in which the fatty acid was attached via an amide linkage at the N terminus were not encephalitogenic, and they induced greater proportions of CD8+ cells in initial in vitro stimulation. Therefore, the lability and the site of the linkage between the peptide and fatty acid may be important for induction of encephalitogenic CD4+ T cells. These results suggest that immune responses induced by endogenous thioacylated lipopeptides may contribute to the immunopathogenesis of chronic experimental demyelinating diseases and multiple sclerosis.

I want you to check the document below with ID 11675392 and I want your output to have only a comma separated list of terms that you consider protein complexes based on the rules. If you think something is a protein complex but consists of affixes (e.g. - or /) and markers, even when they are part of the same syntactic word with a Complex name, as long as there is a separating nonalphanumeric character, do not report them. And remember to check if the terms you want to report are proteins or families.: Phosphoisoprenoids modulate association of Rab geranylgeranyltransferase with REP-1. Rab geranylgeranyltransferase (RabGGTase or GGTase-II) catalyzes the post-translational prenylation of Rab proteins. Rab proteins are recognized as substrates only when they are complexed to Rab Escort Protein (REP). The classical model of prenylation complex assembly assumes initial formation of the Rab.REP binary complex, which subsequently binds to RabGGTase loaded with the isoprenoid donor geranylgeranyl pyrophosphate (GGpp). We demonstrate here that REP-1 can also associate with RabGGTase in the absence of Rab protein and that this interaction is dramatically strengthened by the presence of phosphoisoprenoids such as GGpp. The GGpp-dependent interaction between RabGGTase and REP-1 was observed using affinity precipitations and gel filtration and was quantitated on the basis of fluorescence assays. In the presence of GGpp, REP-1 binds to RabGGTase with a K(d) value of approximately 10 nm, while in its absence the affinity between the two proteins is in the micromolar range. We further demonstrate that binding of Rab7 to the RabGGTase.GGpp.REP-1 complex occurs without prior dissociation of REP-1. Analysis of binding and prenylation rate constants indicate that the RabGGTase.GGpp.REP-1 complex can function as a kinetically competent intermediate of the prenylation reaction. We conclude that, depending on the prevailing concentrations, binding of REP-1 to RabGGTase in the presence of GGpp may serve as an alternative pathway for the assembly of the prenylation machinery in vivo. Implications of these findings for the role of REP-1 in the prenylation reaction are discussed.

I want you to check the document below with ID 11832247 and I want your output to have only a comma separated list of terms that you consider protein complexes based on the rules. If you think something is a protein complex but consists of affixes (e.g. - or /) and markers, even when they are part of the same syntactic word with a Complex name, as long as there is a separating nonalphanumeric character, do not report them. And remember to check if the terms you want to report are proteins or families.: Direct interaction between Rab3b and the polymeric immunoglobulin receptor controls ligand-stimulated transcytosis in epithelial cells. We have examined the role of rab3b in epithelial cells. In MDCK cells, rab3b localizes to vesicular structures containing the polymeric immunoglobulin receptor (pIgR) and located subjacent to the apical surface. We found that GTP-bound rab3b directly interacts with the cytoplasmic domain of pIgR. Binding of dIgA to pIgR causes a dissociation of the interaction with rab3b, a process that requires dIgA-mediated signaling, Arg657 in the cytoplasmic domain of pIgR, and possibly GTP hydrolysis by rab3b. Binding of dIgA to pIgR at the basolateral surface stimulates subsequent transcytosis to the apical surface. Overexpression of GTP-locked rab3b inhibits dIgA-stimulated transcytosis. Together, our data demonstrate that a rab protein can bind directly to a specific cargo protein and thereby control its trafficking.

I want you to check the document below with ID 11917125 and I want your output to have only a comma separated list of terms that you consider protein complexes based on the rules. If you think something is a protein complex but consists of affixes (e.g. - or /) and markers, even when they are part of the same syntactic word with a Complex name, as long as there is a separating nonalphanumeric character, do not report them. And remember to check if the terms you want to report are proteins or families.: Human receptors for sweet and umami taste. The three members of the T1R class of taste-specific G protein-coupled receptors have been hypothesized to function in combination as heterodimeric sweet taste receptors. Here we show that human T1R2/T1R3 recognizes diverse natural and synthetic sweeteners. In contrast, human T1R1/T1R3 responds to the umami taste stimulus l-glutamate, and this response is enhanced by 5'-ribonucleotides, a hallmark of umami taste. The ligand specificities of rat T1R2/T1R3 and T1R1/T1R3 correspond to those of their human counterparts. These findings implicate the T1Rs in umami taste and suggest that sweet and umami taste receptors share a common subunit.

I want you to check the document below with ID 12022229 and I want your output to have only a comma separated list of terms that you consider protein complexes based on the rules. If you think something is a protein complex but consists of affixes (e.g. - or /) and markers, even when they are part of the same syntactic word with a Complex name, as long as there is a separating nonalphanumeric character, do not report them. And remember to check if the terms you want to report are proteins or families.: Yeast Pescadillo is required for multiple activities during 60S ribosomal subunit synthesis. The Pescadillo protein was identified via a developmental defect and implicated in cell cycle progression. Here we report that human Pescadillo and its yeast homolog (Yph1p or Nop7p) are localized to the nucleolus. Depletion of Nop7p leads to nuclear accumulation of pre-60S particles, indicating a defect in subunit export, and it interacts genetically with a tagged form of the ribosomal protein Rpl25p, consistent with a role in subunit assembly. Two pre-rRNA processing pathways generate alternative forms of the 5.8S rRNA, designated 5.8S(L) and 5.8Ss. In cells depleted for Nop7p, the 27SA3 pre-rRNA accumulated, whereas later processing intermediates and the mature 5.8Ss rRNA were depleted. Less depletion was seen for the 5.8S(L) pathway. TAP-tagged Nop7p coprecipitated precursors to both 5.8S(L) and 5.8Ss but not the mature rRNAs. We conclude that Nop7p is required for efficient exonucleolytic processing of the 27SA3 pre-rRNA and has additional functions in 60S subunit assembly and transport. Nop7p is a component of at least three different pre-60S particles, and we propose that it carries out distinct functions in each of these complexes.

I want you to check the document below with ID 14592967 and I want your output to have only a comma separated list of terms that you consider protein complexes based on the rules. If you think something is a protein complex but consists of affixes (e.g. - or /) and markers, even when they are part of the same syntactic word with a Complex name, as long as there is a separating nonalphanumeric character, do not report them. And remember to check if the terms you want to report are proteins or families.: Cytosolic HSP90 associates with and modulates the Arabidopsis RPM1 disease resistance protein. The Arabidopsis protein RPM1 activates disease resistance in response to Pseudomonas syringae proteins targeted to the inside of the host cell via the bacterial type III delivery system. We demonstrate that specific mutations in the ATP-binding domain of a single Arabidopsis cytosolic HSP90 isoform compromise RPM1 function. These mutations do not affect the function of related disease resistance proteins. RPM1 associates with HSP90 in plant cells. The Arabidopsis proteins RAR1 and SGT1 are required for the action of many R proteins, and display some structural similarity to HSP90 co-chaperones. Each associates with HSP90 in plant cells. Our data suggest that (i) RPM1 is an HSP90 client protein; and (ii) RAR1 and SGT1 may function independently as HSP90 cofactors. Dynamic interactions among these proteins can regulate RPM1 stability and function, perhaps similarly to the formation and regulation of animal steroid receptor complexes.

I want you to check the document below with ID 15220471 and I want your output to have only a comma separated list of terms that you consider protein complexes based on the rules. If you think something is a protein complex but consists of affixes (e.g. - or /) and markers, even when they are part of the same syntactic word with a Complex name, as long as there is a separating nonalphanumeric character, do not report them. And remember to check if the terms you want to report are proteins or families.: Silent information regulator 2 potentiates Foxo1-mediated transcription through its deacetylase activity. Longevity regulatory genes include the Forkhead transcription factor FOXO and the NAD-dependent histone deacetylase silent information regulator 2 (Sir2). Genetic studies demonstrate that Sir2 acts to extend lifespan in Caenorhabditis elegans upstream of DAF-16, a member of the FOXO family, in the insulin-like signaling pathway. However, the molecular mechanisms underlying the requirement of DAF-16 activity in Sir2-mediated longevity remain unknown. Here we show that reversible acetylation of Foxo1 (also known as FKHR), the mouse DAF-16 ortholog, modulates its transactivation function. cAMP-response element-binding protein (CREB)-binding protein binds and acetylates Foxo1 at the K242, K245, and K262 residues, the modification of which is involved in the attenuation of Foxo1 as a transcription factor. Conversely, Sir2 binds and deacetylates Foxo1 at residues acetylated by cAMP-response element-binding protein-binding protein. Sir2 is recruited to insulin response sequence-containing promoter and increases the expression of manganese superoxide dismutase and p27(kip1) in a deacetylase-activity-dependent manner. Our findings establish Foxo1 as a direct and functional target for Sir2 in mammalian systems.

I want you to check the document below with ID 16274718 and I want your output to have only a comma separated list of terms that you consider protein complexes based on the rules. If you think something is a protein complex but consists of affixes (e.g. - or /) and markers, even when they are part of the same syntactic word with a Complex name, as long as there is a separating nonalphanumeric character, do not report them. And remember to check if the terms you want to report are proteins or families.: S-palmitoylation modulates estrogen receptor alpha localization and functions. 17beta-Estradiol (E2) acts as a chemical messenger in target tissues inducing both slow nuclear and rapid extra-nuclear responses. E2 binds to its cognate nuclear receptors (ER) resulting in the activation of target gene transcription in the nucleus. In addition to these genomic effects, E2 modulates cell functions through rapid non-genomic actions. Stimulation of G-proteins, Ca(2+) influx, inositol phosphate generation as well as phospholipase C, ERK/MAPK, and PI3K/AKT activation all occur within seconds to minutes after E2 binding to a small population of ERalpha located at the plasma membrane. The great impact of these rapid signals on cell physiology renders central the knowledge of the structural bases and mechanisms that mediate extra-nuclear signaling by E2. Several laboratories, including our own, have recently elucidated the structural requirements for localization and function of plasma membrane ERalpha. This review summarizes the molecular mechanisms of E2-induced rapid non-genomic actions relevant for cell functions, highlighting the role of lipid modification (i.e., palmitoylation) in the ERalpha localization to and residence at the plasma membrane.

I want you to check the document below with ID 17911105 and I want your output to have only a comma separated list of terms that you consider protein complexes based on the rules. If you think something is a protein complex but consists of affixes (e.g. - or /) and markers, even when they are part of the same syntactic word with a Complex name, as long as there is a separating nonalphanumeric character, do not report them. And remember to check if the terms you want to report are proteins or families.: Active gamma-secretase complexes contain only one of each component. Gamma-secretase is an intramembrane aspartyl protease complex that cleaves type I integral membrane proteins, including the amyloid beta-protein precursor and the Notch receptor, and is composed of presenilin, Pen-2, nicastrin, and Aph-1. Although all four of these membrane proteins are essential for assembly and activity, the stoichiometry of the complex is unknown, with the number of presenilin molecules present being especially controversial. Here we analyze functional gamma-secretase complexes, isolated by immunoprecipitation from solubilized membrane fractions and able to produce amyloid beta-peptides and amyloid beta-protein precursor intracellular domain. We show that the active isolated protease contains only one presenilin per complex, which excludes certain models of the active site that require aspartate dyads formed between two presenilin molecules. We also quantified components in the isolated complexes by Western blot using protein standards and found that the amounts of Pen-2 and nicastrin were the same as that of presenilin. Moreover, we found that one Aph-1 was not co-immunoprecipitated with another in active complexes, evidence that Aph-1 is likewise present as a monomer. Taken together, these results demonstrate that the stoichiometry of gamma-components presenilin:Pen-2:nicastrin:Aph-1 is 1:1:1:1.

I want you to check the document below with ID 18253500 and I want your output to have only a comma separated list of terms that you consider protein complexes based on the rules. If you think something is a protein complex but consists of affixes (e.g. - or /) and markers, even when they are part of the same syntactic word with a Complex name, as long as there is a separating nonalphanumeric character, do not report them. And remember to check if the terms you want to report are proteins or families.: Fine mapping of posttranslational modifications of the linker histone H1 from Drosophila melanogaster. The linker histone H1 binds to the DNA in between adjacent nucleosomes and contributes to chromatin organization and transcriptional control. It is known that H1 carries diverse posttranslational modifications (PTMs), including phosphorylation, lysine methylation and ADP-ribosylation. Their biological functions, however, remain largely unclear. This is in part due to the fact that most of the studies have been performed in organisms that have several H1 variants, which complicates the analyses. We have chosen Drosophila melanogaster, a model organism, which has a single H1 variant, to approach the study of the role of H1 PTMs during embryonic development. Mass spectrometry mapping of the entire sequence of the protein showed phosphorylation only in the ten N-terminal amino acids, mostly at S10. For the first time, changes in the PTMs of a linker H1 during the development of a multicellular organism are reported. The abundance of H1 monophosphorylated at S10 decreases as the embryos age, which suggests that this PTM is related to cell cycle progression and/or cell differentiation. Additionally, we have found a polymorphism in the protein sequence that can be mistaken with lysine methylation if the analysis is not rigorous.

I want you to check the document below with ID 18342605 and I want your output to have only a comma separated list of terms that you consider protein complexes based on the rules. If you think something is a protein complex but consists of affixes (e.g. - or /) and markers, even when they are part of the same syntactic word with a Complex name, as long as there is a separating nonalphanumeric character, do not report them. And remember to check if the terms you want to report are proteins or families.: The Glc7 phosphatase subunit of the cleavage and polyadenylation factor is essential for transcription termination on snoRNA genes. Glc7, the yeast protein phosphatase 1, is a component of the cleavage and polyadenylation factor (CPF). Here we show that downregulation of Glc7, or its dissociation from CPF in the absence of CPF subunits Ref2 or Swd2, results in similar snoRNA termination defects. Overexpressing a C-terminal fragment of Sen1, a superfamily I helicase required for snoRNA termination, suppresses the growth and termination defects associated with loss of Swd2 or Ref2, but not Glc7. Suppression by Sen1 requires nuclear localization and direct interaction with Glc7, which can dephosphorylate Sen1 in vitro. The suppressing fragment, and in a similar manner full-length Sen1, copurifies with the snoRNA termination factors Nrd1 and Nab3, suggesting loss of Glc7 from CPF can be compensated by recruiting Glc7 to Nrd1-Nab3 through Sen1. Swd2 is also a subunit of the Set1c histone H3K4 methyltransferase complex and is required for its stability and optimal methyltransferase activity.

I want you to check the document below with ID 19608646 and I want your output to have only a comma separated list of terms that you consider protein complexes based on the rules. If you think something is a protein complex but consists of affixes (e.g. - or /) and markers, even when they are part of the same syntactic word with a Complex name, as long as there is a separating nonalphanumeric character, do not report them. And remember to check if the terms you want to report are proteins or families.: The trafficking/interaction of eNOS and caveolin-1 induced by insulin modulates endothelial nitric oxide production. Endothelial nitric oxide synthase (eNOS) activity is tightly regulated by posttranscriptional modification and its subcellular localization. Here we examined whether insulin modulates nitric oxide (NO) production by regulating eNOS subcellular localization. We used confocal microscopy and immunoblots to examine the time course for 1) subcellular targeting/association of eNOS and caveolin-1 (CAV-1); 2) eNOS Ser(1179) phosphorylation; and 3) NO production in cultured bovine aorta endothelial cells. Serum starvation increased eNOS/CAV-1 localization to the perinuclear region. Adding insulin provoked their prompt translocation to and association at the plasma membrane (PM). Specific monoclonal antibodies against either CAV-1 or eNOS coimmunoprecipitated the other from bovine aorta endothelial cell membrane extracts, and insulin increased this interaction. Insulin stimulated NO production transiently despite a persistent eNOS Ser(1179) phosphorylation. The decline of NO production correlated temporally to insulin-induced translocation of eNOS and CAV-1 to PM. Knockdown of CAV-1 expression with a specific small interfering RNA duplex resulted in eNOS redistributing to the perinuclear region and nearly doubled insulin-induced NO production. Inhibition of phosphatidylinositol 3-kinase activity with wortmannin not only significantly inhibited insulin-induced translocation of eNOS and CAV-1 to PM but also blocked insulin-induced interaction of CAV-1 with eNOS at PM. Insulin increased incorporation of [(3)H]palmitic acid into eNOS immunoprecipitates by approximately 140%. Insulin-induced translocation of eNOS and CAV-1 to PM was palmitoylation dependent. Inhibiting eNOS and CAV-1 palmitoylation enhanced the NO production while blocking the translocation of eNOS and CAV-1 to PM induced by insulin. These data show that insulin acutely regulates eNOS and CAV-1 trafficking to PM of vascular endothelial cells where their interaction can regulate eNOS activity.

I want you to check the document below with ID 21884581_16 and I want your output to have only a comma separated list of terms that you consider protein complexes based on the rules. If you think something is a protein complex but consists of affixes (e.g. - or /) and markers, even when they are part of the same syntactic word with a Complex name, as long as there is a separating nonalphanumeric character, do not report them. And remember to check if the terms you want to report are proteins or families.:    At 24 h before transfection, cells (2 x 105-aliquots) were plated in 6-well plates and transfected with increasing amounts of B7.1-YPet-, B7-H1-YPet- or IR-YPet-expressing plasmids (10 to 500 ng/well), and constant amounts (10 ng-aliquots) of plasmid expressing B7-H1-Rluc8 or B7.1-Rluc8 fusion protein. 48 h later, cells were collected and washed twice with PBS, and aliquots were placed in 384-well plates. Coelenterazine H substrate was added at a final molarity of 5 muM, and BRET was measured immediately. To analyze the effect of the penton, cells were incubated for 5 min without or with penton protein solution at 0.33, 0.66 or 132 ng/muL, followed by Coelenterazine H addition and BRET measurement. BRET was monitored using a lumino/fluorometer (Mithras; Berthold Technologies, France), allowing for the sequential integration of luminescence with two filter settings (Rluc filter, 485 +/- 10 nm; YFP filter, 530 +/- 12.5 nm). The emission signal values obtained at 530 nm were divided by the emission signal values obtained at 485 nm. The BRET ratio was calculated by dividing the emission signal value obtained with coexpressed donor and acceptor by that obtained with the donor protein expressed alone. Data from at least three independent experiments were averaged, and results expressed as milliBRET (mBRET), corresponding to the BRET ratio multiplied by 1,000. Donor saturation curves were determined as previously described.

I want you to check the document below with ID 22723847_14 and I want your output to have only a comma separated list of terms that you consider protein complexes based on the rules. If you think something is a protein complex but consists of affixes (e.g. - or /) and markers, even when they are part of the same syntactic word with a Complex name, as long as there is a separating nonalphanumeric character, do not report them. And remember to check if the terms you want to report are proteins or families.:    Given the role of Ubp9, Ubp13 and Duf1 for normal mitochondria function, we first checked the localization of these three proteins. We found that these three proteins had similar distributions: GFP-tagged proteins were found mostly in the cytoplasm, in various growth conditions (data not shown), as described in databases for Ubp9 and Duf1. Biochemical fractionation of chromosomal-encoded HA-tagged proteins indicated that, in addition to the cytoplasmic soluble fraction, these proteins also display a membrane-bound fraction, possibly associated with mitochondria (Supplemental Fig. S2 B-C).

I want you to check the document below with ID 23028367 and I want your output to have only a comma separated list of terms that you consider protein complexes based on the rules. If you think something is a protein complex but consists of affixes (e.g. - or /) and markers, even when they are part of the same syntactic word with a Complex name, as long as there is a separating nonalphanumeric character, do not report them. And remember to check if the terms you want to report are proteins or families.:    Rif2 promotes a telomere fold-back structure through Rpd3L recruitment in budding yeast.  Using a genome-wide screening approach, we have established the genetic requirements for proper telomere structure in Saccharomyces cerevisiae. We uncovered 112 genes, many of which have not previously been implicated in telomere function, that are required to form a fold-back structure at chromosome ends. Among other biological processes, lysine deacetylation, through the Rpd3L, Rpd3S, and Hda1 complexes, emerged as being a critical regulator of telomere structure. The telomeric-bound protein, Rif2, was also found to promote a telomere fold-back through the recruitment of Rpd3L to telomeres. In the absence of Rpd3 function, telomeres have an increased susceptibility to nucleolytic degradation, telomere loss, and the initiation of premature senescence, suggesting that an Rpd3-mediated structure may have protective functions. Together these data reveal that multiple genetic pathways may directly or indirectly impinge on telomere structure, thus broadening the potential targets available to manipulate telomere function.

I want you to check the document below with ID 23427262_17 and I want your output to have only a comma separated list of terms that you consider protein complexes based on the rules. If you think something is a protein complex but consists of affixes (e.g. - or /) and markers, even when they are part of the same syntactic word with a Complex name, as long as there is a separating nonalphanumeric character, do not report them. And remember to check if the terms you want to report are proteins or families.: Our previous study showed that the major role of the loop lies in the recruitment of the microtubule-associated protein Dis1, a founding member of the TOG/XMAP215 microtubule-associated protein family, and in the ndc80-21 mutant (L405P), Dis1 delocalized from the mitotic kinetochore. Accordingly, we examined Dis1 localization in the ndc80-NH12 mutant and quantified its intensity. Intriguingly, the signal intensities of Dis1 at the mitotic kinetochore were indistinguishable from those of wild-type cells (Figure 4A).

I want you to check the document below with ID 26320581_12 and I want your output to have only a comma separated list of terms that you consider protein complexes based on the rules. If you think something is a protein complex but consists of affixes (e.g. - or /) and markers, even when they are part of the same syntactic word with a Complex name, as long as there is a separating nonalphanumeric character, do not report them. And remember to check if the terms you want to report are proteins or families.: The catalytic domain of MLL4(tag) adopts the canonical beta-fold structure observed for other SET domains, and closely resembles that of MLL1 (Figures 3C and 3D). The N-flanking region of MLL4 forms an extended helical structure, and is markedly different from the extended loop observed for MLL1. Conserved features include the cofactor binding site, which is located in a surface pocket created by the intersection of the SET-N, SET-C, and postSET domains. The position of the substrate binding channel is located between the SET-I and postSET domains, and is indicated in Figure 3C. The C-terminal tag immediately follows the conserved Cys4Zn cluster and enters the substrate binding channel before turning and passing over the cofactor. The tag was required for crystallization but resulted in a catalytically dead construct (Figure 3B), and its impact on the MLL4 SET domain structure and activity is discussed in detail in Figure S3. An equivalent MLL4 construct prepared with an N-terminal tag, like the MLL4(WIN) construct, had high intrinsic activity (Figure 3B), indicating that the determinants of this intrinsic activity are contained within this region of the SET domain. The lack of methyltransferase activity observed with the C-terminally tagged construct can be attributed to partial blocking of the substrate binding channel and interactions with the cofactor.

I want you to check the document below with ID 7667268 and I want your output to have only a comma separated list of terms that you consider protein complexes based on the rules. If you think something is a protein complex but consists of affixes (e.g. - or /) and markers, even when they are part of the same syntactic word with a Complex name, as long as there is a separating nonalphanumeric character, do not report them. And remember to check if the terms you want to report are proteins or families.: Evolutionary conservation of human TATA-binding-polypeptide-associated factors TAFII31 and TAFII80 and interactions of TAFII80 with other TAFs and with general transcription factors. Human transcription initiation factor TFIID is composed of the TATA-binding polypeptide (TBP) and at least 13 TBP-associated factors (TAFs) that collectively or individually are involved in activator-dependent transcription. To investigate protein-protein interactions involved in TFIID assembly and in TAF-mediated activator functions, we have cloned and expressed cDNAs encoding human TAFII80 and TAFII31. Coimmunoprecipitation assays showed that TAFII80 interacted with TAFII250, TAFII31, TAFII20, and TBP, but not with TAFII55. Similar assays showed that TAFII80 interacted with TFIIE alpha and with TFIIF alpha (RAP74) but not with TFIIB, TFIIE beta, or TFIIF beta (RAP30). Further studies with TAFII80 mutations revealed three distinct interaction domains which fall within regions conserved in human TAFII80, Drosophila TAFII60, and yeast TAFII60. The N terminus of TAFII80 (residues 1-100) interacts with both TAFII31 and TAFII20, while two C-terminal regions are involved, respectively, in interactions with TAFII250 and TFIIF alpha (RAP74) (residues 203-276) and with TBP and TFIIE alpha (residues 377-505). The interactions between TAFII80 and general factors TFIIE alpha and TFIIF alpha (RAP74) could be important for recruitment of GTFs during activator-dependent transcription. Because TAFs 80, 31, and 20 show sequence similarities to histones H4, H3, and H2B, as well as some parallel interactions, this subset of TAFs may form a related core structure within TFIID.

I want you to check the document below with ID 7957109 and I want your output to have only a comma separated list of terms that you consider protein complexes based on the rules. If you think something is a protein complex but consists of affixes (e.g. - or /) and markers, even when they are part of the same syntactic word with a Complex name, as long as there is a separating nonalphanumeric character, do not report them. And remember to check if the terms you want to report are proteins or families.: A proteasome inhibitor prevents activation of NF-kappa B and stabilizes a newly phosphorylated form of I kappa B-alpha that is still bound to NF-kappa B. Activation of the inducible transcription factor NF-kappa B involves removal of the inhibitory subunit I kappa B-alpha from a latent cytoplasmic complex. It has been reported that I kappa B-alpha is subject to both phosphorylation and proteolysis in the process of NF-kappa B activation. In this study, we present evidence that the multicatalytic cytosolic protease (proteasome) is involved in the degradation of I kappa B-alpha. Micromolar amounts of the peptide Cbz-Ile-Glu(O-t-Bu)-Ala-leucinal (PSI), a specific inhibitor of the chymotrypsin-like activity of the proteasome, prevented activation of NF-kappa B in response to tumor necrosis factor-alpha (TNF) and okadaic acid (OA) through inhibition of I kappa B-alpha degradation. The m-calpain inhibitor Cbz-Leu-leucinal was ineffective. In the presence of PSI, a newly phosphorylated form of I kappa B-alpha accumulated in TNF- and OA-stimulated cells. However, the covalent modification of I kappa B-alpha was not sufficient for activation of NF-kappa B: no substantial NF-kappa B DNA binding activity appeared in cells because the newly phosphorylated form of I kappa B-alpha was still tightly bound to p65 NF-kappa B. Pyrrolidinedithiocarbamate, an antioxidant inhibitor of NF-kappa B activation which did not interfere with proteasome activities, prevented de novo phosphorylation of I kappa B-alpha as well as its subsequent degradation. This suggests that phosphorylation of I kappa B-alpha is equally necessary for the activation of NF-kappa B.(ABSTRACT TRUNCATED AT 250 WORDS)

I want you to check the document below with ID 8506307 and I want your output to have only a comma separated list of terms that you consider protein complexes based on the rules. If you think something is a protein complex but consists of affixes (e.g. - or /) and markers, even when they are part of the same syntactic word with a Complex name, as long as there is a separating nonalphanumeric character, do not report them. And remember to check if the terms you want to report are proteins or families.: In vitro and in vivo functional characterization of bovine vitamin K-dependent gamma-carboxylase expressed in Chinese hamster ovary cells. Coagulation factor IX is a serine protease for which high-level expression of biologically active protein in heterologous cells is limited due to inefficient proteolytic removal of the propeptide as well as vitamin K-dependent carboxylation of multiple amino-terminal glutamic acid residues. We have overexpressed the vitamin K-dependent gamma-carboxylase cDNA and monitored its ability to improve factor IX processing in Chinese hamster ovary (CHO) cells. From amino acid sequence analysis of bovine liver vitamin K-dependent gamma-carboxylase, degenerate oligonucleotides were used to isolate a 3.5-kbp bovine cDNA that encoded a 758-residue open reading frame. Expression of the cDNA in COS-1 and CHO cells yielded 17- and 16-fold increases in the in vitro gamma-carboxylase activity of microsomal preparations, respectively. Anti-serum raised against a predicted peptide sequence reacted with a 94-kDa polypeptide in the partially purified bovine liver preparation as well as in stably transfected CHO cells. The amount of antibody reactivity correlated with the increased ability to carboxylate a peptide substrate in vitro. These results strongly support the conclusion that the cDNA encodes the vitamin K-dependent gamma-carboxylase. Transient transfection of the gamma-carboxylase expression vector into factor IX-expressing CHO cells did not improve the specific procoagulant activity of secreted factor IX. In contrast, transfection of an expression vector encoding the propeptide processing enzyme PACE (paired basic amino acid cleaving enzyme) did improve the specific activity of secreted factor IX by 3-fold. These results demonstrate that the ability of CHO cells to modify glutamic acid residues to gamma-carboxyglutamic acid in secreted factor IX is not limited by the expression of the vitamin K-dependent gamma-carboxylase alone.

I want you to check the document below with ID 9616112 and I want your output to have only a comma separated list of terms that you consider protein complexes based on the rules. If you think something is a protein complex but consists of affixes (e.g. - or /) and markers, even when they are part of the same syntactic word with a Complex name, as long as there is a separating nonalphanumeric character, do not report them. And remember to check if the terms you want to report are proteins or families.: Role of the CLOCK protein in the mammalian circadian mechanism. The mouse Clock gene encodes a bHLH-PAS protein that regulates circadian rhythms and is related to transcription factors that act as heterodimers. Potential partners of CLOCK were isolated in a two-hybrid screen, and one, BMAL1, was coexpressed with CLOCK and PER1 at known circadian clock sites in brain and retina. CLOCK-BMAL1 heterodimers activated transcription from E-box elements, a type of transcription factor-binding site, found adjacent to the mouse per1 gene and from an identical E-box known to be important for per gene expression in Drosophila. Mutant CLOCK from the dominant-negative Clock allele and BMAL1 formed heterodimers that bound DNA but failed to activate transcription. Thus, CLOCK-BMAL1 heterodimers appear to drive the positive component of per transcriptional oscillations, which are thought to underlie circadian rhythmicity.

I want you to check the document below with ID 9659924 and I want your output to have only a comma separated list of terms that you consider protein complexes based on the rules. If you think something is a protein complex but consists of affixes (e.g. - or /) and markers, even when they are part of the same syntactic word with a Complex name, as long as there is a separating nonalphanumeric character, do not report them. And remember to check if the terms you want to report are proteins or families.: Recruitment of CBP/p300 by the IFN beta enhanceosome is required for synergistic activation of transcription. Transcriptional activation of the IFN beta gene in response to virus infection requires the assembly of an enhanceosome, consisting of the transcriptional activators NF-kappa B, IRF1, ATF2/c-Jun, and the architectural protein HMG I(Y). The level of transcription generated by all of these activators is greater than the sum of the levels generated by individual factors, a phenomenon designated transcriptional synergy. We demonstrate that this synergy, in the context of the enhanceosome, requires a new protein-protein interaction domain in the p65 subunit of NF-kappa B. Transcriptional synergy requires recruitment of the CBP/p300 coactivator to the enhanceosome, via a new activating surface assembled from the novel p65 domain and the activation domains of all of the activators. Deletion, substitution, or rearrangement of any one of the activation domains in the context of the enhanceosome decreases both recruitment of CBP and transcriptional synergy.

### Supplementary Section 2: CoNECO GPT full - Instructions

The “CoNECo GPT - full” available through https://chat.openai.com/g/g-1uV7nfJTA-coneco-gpt-full is a custom ChatGPT that was generated with the following set of instructions corresponding to the annotation guidelines provided to human annotators. There is a limit of 8000 characters for instructions and for that reason the GO term names were removed from the list in the bottom of the instructions:

You are a helpful assistant. Your task is to identify protein-containing complexes in the CoNECo corpus using the following annotation guidelines:
- Entities that fall within the scope of the Gene Ontology term protein-containing complex are the target of named entity annotations for CoNECo. Excerpt taken from Gene Ontology: A protein complex in this context is meant as a stable set of interacting proteins which can be co-purified by an acceptable method, and where the complex has been shown to exist as an isolated, functional unit in vivo. Acceptable experimental methods include stringent protein purification followed by detection of protein interaction. The following methods should be considered non-acceptable: simple immunoprecipitation, pull-down experiments from cell extracts without further purification, colocalization and 2-hybrid screening. Interactions that should not be captured as protein complexes include: 1) enzyme/substrate, receptor/ligand or any similar transient interactions, unless these are a critical part of the complex assembly or are required e.g. for the receptor to be functional; 2) proteins associated in a pull-down/co-immunoprecipitation assay with no functional link or any evidence that this is a defined biological entity rather than a loose-affinity complex; 3) any complex where the only evidence is based on genetic interaction data; 4) partial complexes, where some subunits (e.g. transmembrane ones) cannot be expressed as recombinant proteins and are excluded from experiments (in this case, independent evidence is necessary to find out the composition of the full complex, if known). Interactions that may be captured as protein complexes include: 1) enzyme/substrate or receptor/ligand if the complex can only assemble and become functional in the presence of both classes of subunits; 2) complexes where one of the members has not been shown to be physically linked to the other(s), but is a homologue of, and has the same functionality as, a protein that has been experimentally demonstrated to form a complex with the other member(s); 3) complexes whose existence is accepted based on localization and pharmacological studies, but for which experimental evidence is not yet available for the complex as a whole.
- If a term is found in Gene ontology but it is NOT a protein-containing complex, then it will NOT be considered a Complex in this effort (e.g. ribosome).
- Gene Ontology-Cellular Component (GO-CC) and specifically GO terms under protein-containing complex will be the target for Named Entity Normalization.
- If a protein-containing complex can be identified as such by the annotators, but there is no entry in Gene Ontology corresponding to the entity then it will receive an annotation as a named entity (NE), but will not have a normalization to GO-CC.
- The first resource that is trusted to resolve issues is Gene Ontology. If there is still not enough information there, inconsistencies will be resolved using Complex Portal, Reactome, CORUM, and as a last result the literature.
- For each mention of a protein-containing complex (Complex hereafter) name, the annotation aims to mark the minimal span containing the full name of the entity mentioned in the text so that the marked span starts and ends on a boundary between an alphanumeric string and a non-alphanumeric character (e.g. space or hyphen). The following provides examples and guidelines for exceptional cases.
- Modifiers and head words that are not part of the name are excluded from the annotated span. This includes the word “complex” which should not be annotated as part of the entity. For example in "human NFkappaB complex", only "NFkappaB" is annotated and in "phosphorylated RNA polymerase II" only "RNA polymerase II" is annotated.

- Affixes and markers are similarly excluded from the annotation span even when they are part of the same syntactic word with a Complex name as long as there is a separating nonalphanumeric character, for example in "phospho-RNA polymerase II" only "RNA polymerase II" is annotated.
- However, annotation boundaries must coincide with the boundary between an alphanumeric and a non-alphanumeric character. In cases where a Complex name is written with one of the following regular affixes without such a boundary, the affix is included in the annotated span: i) species identifier, e.g. h for human or m for mouse, ii) p for phosphorylated, iii) wt for wild-type, iv) si for small interfering RNA, v) sh for small/short hairpin RNA, vi) anti for antibody. For example in "the human complex hNFkappaB, and the murine complex mNFkappaB", "hNFkappa B" and "mNFkappaB" are annotated and in "pRNApolII (phosphor RNA polymerase II)", "pRNApolII" and "RNA polymerase II" are annotated.
- When a protein-containing complex name coincides with the name of a GGP or the GGPs comprising it or the name of a Protein Family, separated by ANY punctuation, annotation of the Protein or Protein Family named entities is preferred over annotation of the protein-containing complex entity. For example in "ERMES complex (otherwise known as Mdm10/Mdm12/Mmm1 complex)" only "ERMES" is annotated and "Mdm10/Mdm12/Mmm1" are not annotated as a single Complex entity, as they constitute three separate Protein NEs
- Two notable exceptions are "Arp2/3" and "SWI/SNF" where a single Protein-containing complex named entity is annotated instead.
- Annotations should be applied to all variants of a Complex name. For example "NF kappaB", "NF-kappaB", "NFkappaB" should all be marked as Protein-containing complex.
- PDB identifiers (for example "3BMP", "4BQ6") will NOT be annotated as Protein-containing complex even if when checked against PDB they correspond to one.
- List of GO terms which are protein-containing complex in GO, that will NOT be annotated as Complex as they consitute groups of complexes.

* GO:1990104
* GO:0140535
* GO:1902494
* GO:0070069
* GO:0031074
* GO:0005875
* GO:0001114
* GO:0150005
* GO:0019907
* GO:0044796
* GO:0098636
* GO:0098635
* GO:0098637
* GO:0043235
* GO:0140368
* GO:1903768
* GO:0098666
* GO:1990563
* GO:0019036
* GO:0098796
* GO:0098803
* GO:1990684
* GO:1990686
* GO:1990685
* GO:0140513
* GO:0035097
* GO:0000109
* GO:0008023
* GO:0005849
* GO:0000152

* GO:0032994
* GO:0060987
* GO:1990777
* GO:0030076
* GO:0098798
* GO:0090665
* GO:1990351
* GO:0005667
* GO:0017053
* GO:0090576
* GO:0090577
* GO:0090578
* GO:1903865
* GO:0000120
* GO:0090575
* GO:0032992
* GO:0046806
* GO:0032300
* GO:0032993
* GO:0031588
* GO:1990234
* GO:0061695
* GO:1902911
* GO:0009365
* GO:1902554
* GO:0000307
* GO:1902554
* GO:1902493
* GO:0031248
* GO:0031501
* GO:0106068
* GO:1902503
* GO:0042575
* GO:0030880
* GO:0005965
* GO:0034708
* GO:1990228
* GO:0016459
* GO:0045298
* GO:0031941
* GO:0005942
* GO:0000786
* GO:0038201
* GO:0034360
* GO:0071256
* GO:1990425
* GO:0034270
* GO:0106003

### Supplementary Table 1: CoNECO GPT full - Results

Below are detailed results of the complexes that this custom version of ChatGPT has detected in the 25 documents provided as prompt. We used this list to count True Positives (TP), False Positives (FP) and False Negatives (FN) for each document and for the total set of documents.

| PubMed ID | TP | FP | FN | detected complexes |
|---|---|---|---|---|
| 10359895 | 0 | 6 | 0 | YY1, Oct1 |
| 10660620 | 0 | 6 | 0 | SHP-1 |
| 18342605 | 4 | 19 | 2 | CPF, Glc7, Ref2, Swd2, Sen1, Nrd1-Nab3, Set1c |
| 19608646 | 0 | 29 | 0 | eNOS, CAV-1 |
| 21884581_16 | 0 | 5 | 0 | B7.1-YPet, B7-H1-YPet, IR-YPet, B7-H1-Rluc8, B7.1-Rluc8 |
| 22723847_14 | 0 | 5 | 0 | Ubp9, Ubp13, Duf1 |
| 23028367 | 5 | 0 | 0 | Rpd3L, Rpd3S, Hda1 |
| 23427262_17 | 0 | 4 | 0 | Dis1 |
| 26320581_12 | 0 | 7 | 0 | MLL4, MLL1 |
| 7667268 | 0 | 27 | 3 | TAFII80, TAFII31, TAFII250, TAFII20, TBP, TFIIE alpha, TFIIF alpha (RAP74) |
| 7957109 | 10 | 10 | 4 | NF-kappa B, I kappa B-alpha |
| 8506307 | 0 | 9 | 0 | gamma-carboxylase, PACE |
| 9616112 | 0 | 2 | 0 | CLOCK-BMAL1 |
| 11331609 | 1 | 3 | 0 | NuRD, HDAC1, HDAC2 |
| 11359852 | 0 | 6 | 0 | PLP |
| 11675392 | 8 | 9 | 3 | RabGGTase, REP-1 |
| 11832247 | 0 | 12 | 0 | rab3b, pIgR |
| 11917125 | 0 | 4 | 0 | T1R2/T1R3, T1R1/T1R3 |
| 12022229 | 0 | 6 | 5 | Nop7p |
| 14592967 | 0 | 17 | 0 | HSP90, RPM1, RAR1, SGT1 |
| 15220471 | 0 | 12 | 0 | Sir2, Foxo1 |
| 16274718 | 0 | 3 | 0 | ERalpha |
| 17911105 | 3 | 16 | 0 | gamma-secretase, presenilin, Pen-2, nicastrin, Aph-1 |
| 18253500 | 0 | 8 | 0 | H1 |
| 9659924 | 1 | 0 | 6 | IFN beta enhanceosome |
| Total | 32 | 225 | 23 | |

### Supplementary Section 3: CoNECO GPT small - Instructions

CoNECo GPT - small, available through this link: https://chat.openai.com/g/g-Ns0dcCn8c-coneco-gpt-small was created by providing the following set of instructions:

You are a helpful assistant. Your task is to identify protein-containing complexes in the CoNECo corpus using the following annotation guidelines:
- Entities that fall within the scope of the Gene Ontology term protein-containing complex are the target of named entity annotations for CoNECo: "A protein complex in this context is meant as a stable set of interacting proteins which can be co-purified by an acceptable method, and where the complex has been shown to exist as an isolated, functional unit in vivo. Acceptable experimental methods include stringent protein purification followed by detection of protein interaction. The following methods should be considered non-acceptable: simple immunoprecipitation, pull-down experiments from cell extracts without further purification, colocalization and 2-hybrid screening. Interactions that should not be captured as protein complexes include: 1) enzyme/substrate, receptor/ligand or any similar transient interactions, unless these are a critical part of the complex assembly or are required e.g. for the receptor to be functional; 2) proteins associated in a pull-down/co-immunoprecipitation assay with no functional link or any evidence that this is a defined biological entity rather than a loose-affinity complex; 3) any complex where the only evidence is based on genetic interaction data; 4) partial complexes, where some subunits (e.g. transmembrane ones) cannot be expressed as recombinant proteins and are excluded from experiments (in this case, independent evidence is necessary to find out the composition of the full complex, if known). Interactions that may be captured as protein complexes include: 1) enzyme/substrate or receptor/ligand if the complex can only assemble and become functional in the presence of both classes of subunits; 2) complexes where one of the members has not been shown to be physically linked to the other(s), but is a homologue of, and has the same functionality as, a protein that has been experimentally demonstrated to form a complex with the other member(s); 3) complexes whose existence is accepted based on localization and pharmacological studies, but for which experimental evidence is not yet available for the complex as a whole."
- If a term is found in Gene ontology but it is NOT a protein-containing complex, then it will NOT be considered a Complex in this effort
- If a protein-containing complex can be identified as such by the annotators, but there is no entry in Gene Ontology corresponding to the entity then it will receive an annotation as a named entity (NE).
- The first resource that is trusted to resolve issues is Gene Ontology. If there is still not enough information there, inconsistencies will be resolved using Complex Portal, Reactome, CORUM, and as a last result the literature.
- For each mention of a protein-containing complex (Complex hereafter) name, the annotation aims to mark the minimal span containing the full name of the entity mentioned in the text so that the marked span starts and ends on a boundary between an alphanumeric string and a non-alphanumeric character (e.g. space or hyphen).
- Modifiers and head words that are not part of the name are excluded from the annotated span. This includes the word "complex" which should not be annotated as part of the entity.
- Affixes and markers are similarly excluded from the annotation span even when they are part of the same syntactic word with a Complex name as long as there is a separating nonalphanumeric character.
- However, annotation boundaries must coincide with the boundary between an alphanumeric and a non-alphanumeric character.
- When a protein-containing complex name coincides with the name of a Protein or the Proteins comprising it or the name of a Protein Family, separated by ANY punctuation, annotation of the Protein or Protein Family named entities is preferred over annotation of the protein-containing complex entity.
- Two notable exceptions are "Arp2/3" and "SWI/SNF" where a single Protein-containing complex named entity is annotated instead.

- Annotations should be applied to all variants of a Complex name.
- PDB identifiers will NOT be annotated as Protein-containing complex.
- GO terms which are protein-containing complex in GO, but consitute groups of complexes, will not be annotated as Complex. This includes GO terms the definition of which starts with the phrase: "A protein complex which is capable of...".

### Supplementary Table 2: CoNECO GPT small - Results

Below are detailed results of the complexes that this custom version of ChatGPT has detected in the 25 documents provided as prompt. We once again used this list to count TP, FP and FN for each document and for the total set of documents.

| PubMed ID | TP | FP | FN | detected complexes |
|---|---|---|---|---|
| 10359895 | 0 | 6 | 0 | YY1, Oct1 |
| 10660620 | 0 | 0 | 0 | - |
| 18342605 | 4 | 2 | 0 | CPF, Nrd1-Nab3, Set1c |
| 19608646 | 0 | 29 | 0 | eNOS, CAV-1 |
| 21884581_16 | 0 | 5 | 0 | B7.1-YPet, B7-H1-YPet, IR-YPet, B7-H1-Rluc8, B7.1-Rluc8 |
| 22723847_14 | 0 | 0 | 0 | - |
| 23028367 | 5 | 0 | 0 | Rpd3L, Rpd3S, Hda1 |
| 23427262_17 | 0 | 4 | 0 | Dis1 |
| 26320581_12 | 0 | 7 | 0 | MLL4, MLL1 |
| 7667268 | 3 | 27 | 0 | TFIID,TAFII80,TAFII250,TAFII31,TAFII20,TBP,TFIIE alpha,TFIIF alpha |
| 7957109 | 10 | 10 | 4 | NF-kappa B,I kappa B-alpha |
| 8506307 | 0 | 9 | 0 | gamma-carboxylase, PACE |
| 9616112 | 0 | 2 | 0 | CLOCK-BMAL1 |
| 11331609 | 1 | 2 | 0 | NuRD, SRA |
| 11359852 | 0 | 0 | 0 | - |
| 11675392 | 8 | 7 | 3 | RabGGTase, REP-1, RabGGTase.GGpp.REP-1 |
| 11832247 | 0 | 0 | 0 | - |
| 11917125 | 0 | 4 | 0 | T1R2/T1R3, T1R1/T1R3 |
| 12022229 | 0 | 6 | 5 | Nop7p |
| 14592967 | 0 | 17 | 0 | HSP90, RPM1, RAR1, SGT1 |
| 15220471 | 0 | 12 | 0 | Foxo1,Sir2 |
| 16274718 | 0 | 3 | 0 | ERalpha |
| 17911105 | 3 | 0 | 0 | gamma-secretase |
| 18253500 | 0 | 0 | 0 | - |
| 9659924 | 3 | 5 | 0 | CBP/p300, IFN beta enhanceosome, NF-kappa B, IRF1, ATF2/c-Jun, HMG I(Y) |
| Total | 37 | 157 | 12 | |

### Supplementary Section 4: CoNECO GPT minimal - Instructions

"CoNECo GPT - minimal" is available through the Explore GPTs section of ChatGPT4.0 https://chat.openai.com/g/g-C6Nx12aEL-coneco-gpt-minimal and was generated using the instructions below:

```
Coneco specializes in the identification of protein complexes with unwavering adherence to a set of precise guidelines. It defines protein complexes based on the Gene Ontology term 'protein-containing complex' (GO:0032991) within the cellular component. A protein complex is identified as a stable set of interacting proteins that are co-purifiable and function together in vivo. Methods like stringent protein purification followed by interaction detection are acceptable for validation, but not mere immunoprecipitation or colocalization. Entities not categorized as 'protein-containing complex' in Gene Ontology or not children of GO:0032991, including entities such as ribosomes, are not considered protein complexes by Coneco. In its identification process, Coneco first verifies if an entity is a protein using UniProt, then checks if it's a protein family in InterPro. Entities that are neither proteins nor protein families are then evaluated against Coneco's rules, referencing Gene Ontology, Complex Portal, Reactome, CORUM, and relevant literature. Coneco annotates minimal spans of protein complex names, avoiding non-essential modifiers. Entities coinciding with genes, gene products, or protein families, without evidence of being a functional entity, are not annotated as complexes. Additionally, certain GO-classified protein-containing complexes are excluded from annotation. Coneco has two specific rules to enhance its accuracy: 1) If entities are separated by a dash (-), they are examined and reported separately, with each being individually assessed against the established guidelines. 2) If terms examined are proteins or protein families, they are not reported as protein complexes. Coneco relies on resources like PubMed, GO, InterPro, and UniProt for verification, strictly following these guidelines and definitions.
```

### Supplementary Table 3: CoNECO GPT minimal - Results

The results for the same set of 25 prompts are the following:

| PubMed ID | TP | FP | FN | detected words |
|---|---|---|---|---|
| 10359895 | 0 | 0 | 0 | - |
| 10660620 | 0 | 0 | 0 | - |
| 18342605 | 2 | 2 | 4 | cleavage and polyadenylation factor, Set1c histone H3K4 methyltransferase complex, Nrd1-Nab3 complex |
| 19608646 | 0 | 0 | 0 | - |
| 21884581_16 | 0 | 0 | 0 | - |
| 22723847_14 | 0 | 0 | 0 | - |
| 23028367 | 5 | 0 | 0 | Rpd3L, Rpd3S, Hda1 complexes |
| 23427262_17 | 0 | 0 | 0 | - |
| 26320581_12 | 0 | 0 | 0 | - |
| 7667268 | 3 | 0 | 0 | TFIID |
| 7957109 | 10 | 0 | 4 | NF-kappa B/I kappa B-alpha complex |
| 8506307 | 0 | 0 | 0 | - |
| 9616112 | 0 | 1 | 0 | CLOCK-BMAL1 heterodimers |
| 11331609 | 1 | 0 | 0 | NuRD complex |
| 11359852 | 0 | 0 | 0 | - |
| 11675392 | 2 | 0 | 9 | RabGGTase.GGpp.REP-1 complex |
| 11832247 | 0 | 0 | 0 | - |
| 11917125 | 0 | 4 | 0 | T1R2/T1R3, T1R1/T1R3 |
| 12022229 | 2 | 0 | 3 | pre-60S particles |
| 14592967 | 0 | 0 | 0 | - |
| 15220471 | 0 | 0 | 0 | - |
| 16274718 | 0 | 0 | 0 | - |
| 17911105 | 3 | 0 | 0 | gamma-secretase complex |
| 18253500 | 0 | 0 | 0 | - |
| 9659924 | 1 | 0 | 6 | IFN beta enhanceosome |
| Total | 29 | 7 | 26 | |

Supplementary Table 4: Normalization errors for Jensenlab tagger run on the CoNECo test set.

| PubMed ID | annotated entity | tagged entity | annotated normalization | tagger normalization | comment |
|---|---|---|---|---|---|
| 23499533_3 | cohesin | cohesin loading | GO:0008278 | GO:0032116 | cohesin loading complex is a narrow synonym for SMC complex, and is tagged as the left-longest entity |
| 16209941 | SCF(Skp2) ubiquitin ligase | SCF | GO:0097669 | GO:0019005 | the more general SCF has been tagged and normalized |
| 16209941 | SCF(Skp2) | SCF | GO:0097669 | GO:0019005 | the more general SCF has been tagged and normalized |
| 16209941 | SCF(Skp2) | SCF | GO:0097669 | GO:0019005 | the more general SCF has been tagged and normalized |
| 20427570_13 | Complex III | Complex III | GO:0005750 | GO:0045275 | the more general respiratory complex has been tagged and normalized, instead of the mitochondrial |
| 20427570_13 | complex III | complex III | GO:0005750 | GO:0045275 | the more general respiratory complex has been tagged and normalized, instead of the mitochondrial |
| 21347277_5 | RFC | Ctf18-RFC | GO:0005663 | GO:0031390 | the more specific complex has been tagged and normalized, due to a dictionary error |
| 21347277_5 | RFC | Ctf18-RFC | GO:0005663 | GO:0031390 | the more specific complex has been tagged and normalized, due to a dictionary error |
| 21347277_5 | RFC | Ctf18-RFC | GO:0005663 | GO:0031390 | the more specific complex has been tagged and normalized, due to a dictionary error |
| 15223 | cytochrome c oxidase | cytochrome c oxidase | - | GO:0045277 | annotation error |

### Supplementary Table 5: Error analysis results for Jensenlab tagger run on the CoNECo test set

| PubMed ID | FP/ FN | error count | entity | error type | comment |
|---|---|---|---|---|---|
| 1939122 | FN | 1 | 30 S ribosomal subunits | dictionary error | synonym not in GO |
| 1939122 | FN | 1 | 28 S | dictionary error | synonym not in GO |
| 3453108 | FN | 1 | high density lipoprotein | dictionary error | synonym not in GO |
| 8407894 | FN | 1 | TFIIIB | dictionary error | synonym not in GO |
| 8407894 | FN | 3 | RNA polymerase III | dictionary error | block list |
| 9367446 | FN | 1 | alpha(v)beta1 | dictionary error | synonym not in GO |
| 9857197 | FN | 1 | NPC | dictionary error | block list |
| 10617773 | FN | 2 | collagen | dictionary error | synonym not in GO |
| 10770935 | FN | 3 | DRIP | dictionary error | synonym not in GO |
| 12504026 | FN | 1 | IkappaB | dictionary error | synonym not in GO |
| 12815069 | FN | 2 | TRAIL | dictionary error | synonym not in GO |
| 15257288 | FN | 1 | type I collagen | dictionary error | synonym not in GO |
| 15257288 | FN | 5 | collagen | dictionary error | synonym not in GO |
| 15449939 | FN | 1 | type I human interferon receptor | dictionary error | synonym not in GO |
| 15886116 | FN | 1 | T-cell antigen receptor | dictionary error | synonym not in GO |
| 15886116 | FN | 4 | TCR | dictionary error | block list |
| 16024914 | FN | 1 | HDL | dictionary error | block list |
| 16717095 | FN | 2 | cytosolic chaperonin complex | dictionary error | synonym not in GO |
| 16717095 | FN | 3 | CCT | dictionary error | synonym not in GO |
| 17363904 | FN | 1 | TCR | dictionary error | block list |
| 17532339 | FN | 1 | E1 | dictionary error | block list |
| 17532339 | FN | 3 | PDC | dictionary error | synonym not in GO |
| 17615152 | FN | 1 | activator protein-1 | dictionary error | synonym not in GO |
| 17707230 | FN | 4 | Pol I | dictionary error | synonym not in GO |
| 18391171 | FN | 1 | actomyosin | dictionary error | synonym not in GO |
| 18838202 | FN | 1 | nuclear factor-kappaB | dictionary error | synonym not in GO |
| 18838202 | FN | 1 | tumor necrosis factor (TNF)-related apoptosis-inducing ligand | dictionary error | synonym not in GO |
| 18838202 | FN | 9 | TRAIL | dictionary error | synonym not in GO |

| PubMed ID | FP/ FN | error count | entity | error type | comment |
|---|---|---|---|---|---|
| 19017259 | FN | 1 | tissue factor-activated FVII | dictionary error | synonym not in GO |
| 19503082 | FN | 2 | AMPARs | dictionary error | synonym not in GO |
| 19503082 | FN | 4 | AMPAR | dictionary error | synonym not in GO |
| 19637179_9 | FN | 1 | COX | dictionary error | synonym not in GO |
| 19684574_7 | FN | 1 | RISC | dictionary error | block list |
| 19684574_7 | FN | 7 | miRISC | dictionary error | synonym not in GO |
| 19706605 | FN | 3 | proteosome | dictionary error | synonym not in GO |
| 20427570_13 | FN | 1 | complexes III | dictionary error | synonym not in GO |
| 20427570_13 | FN | 2 | complex IV | dictionary error | synonym not in GO |
| 20427570_13 | FN | 2 | FoF1-ATPase | dictionary error | synonym not in GO |
| 23810392 | FN | 1 | nuclear factor (NF)-kappaB | dictionary error | synonym not in GO |
| 24498436 | FN | 1 | RNA polymerase II | dictionary error | block list |
| 24498436 | FN | 2 | Pol II | dictionary error | synonym not in GO |
| 24498436 | FN | 2 | DDR | dictionary error | term not in GO |
| 24498436 | FN | 3 | Pol V | dictionary error | synonym not in GO |
| 26052893_5 | FN | 1 | ECF transporter | dictionary error | synonym not in GO |
| 26052893_5 | FN | 1 | ECF transporters | dictionary error | synonym not in GO |
| 27350684 | FN | 2 | 3'-end processing | dictionary error | synonym not in GO |
| 28515143_17 | FN | 2 | Mediator | dictionary error | block list |
| 28806752_11 | FN | 1 | IgG | dictionary error | synonym not in GO |
| 20427570_13 | FN | 1 | IV | discontinuous entity | |
| 2462567 | FP | 7 | enolase | ambiguous name | family |
| 2981876 | FP | 4 | p35 | ambiguous name | protein |
| 8641455 | FP | 2 | SNARE | ambiguous name | family |
| 8663127 | FP | 1 | TSC | ambiguous name | other entity |
| 8798525 | FP | 3 | palmitoyltransferase | ambiguous name | family |
| 8932298 | FP | 1 | casein kinase II | ambiguous name | family |
| 9150262 | FP | 3 | global genome repair | ambiguous name | general term |
| 9195882 | FP | 1 | NADPH oxidase | ambiguous name | family |
| 9497260 | FP | 1 | phosphomannomutase | ambiguous name | family |

| PubMed ID | FP/ FN | error count | entity | error type | comment |
|---|---|---|---|---|---|
| 9497260 | FP | 2 | PMM2 | ambiguous name | family |
| 10446169 | FP | 1 | PS1 | ambiguous name | family |
| 10446169 | FP | 4 | presenilins | ambiguous name | family |
| 10666337 | FP | 1 | topoisomerase II | ambiguous name | protein |
| 10763828 | FP | 1 | docking protein | ambiguous name | general term |
| 11121043 | FP | 1 | BMP receptors | ambiguous name | general term |
| 11121043 | FP | 4 | Ski | ambiguous name | protein |
| 11874917 | FP | 2 | nonmuscle myosin | ambiguous name | family |
| 14562105 | FP | 2 | glucose transporter | ambiguous name | general term |
| 14752096 | FP | 13 | AML1 | ambiguous name | protein |
| 16055508 | FP | 1 | Kinesin | ambiguous name | family |
| 18391171 | FP | 1 | capping proteins | ambiguous name | general term |
| 20129058 | FP | 1 | DnaB helicase | ambiguous name | protein |
| 20129058 | FP | 1 | DnaB hexamer | ambiguous name | protein |
| 20129058 | FP | 2 | prepriming complex | ambiguous name | general term |
| 20610737 | FP | 1 | cyclic nucleotide phosphodiesterase | ambiguous name | general term |
| 21807881 | FP | 2 | SNARE | ambiguous name | family |
| 21827752 | FP | 1 | vesicle-tethering complex | ambiguous name | general term |
| 21900752 | FP | 1 | ARF | ambiguous name | family |
| 21900752 | FP | 1 | ARF complex | ambiguous name | family |
| 22194618 | FP | 2 | formate dehydrogenases | ambiguous name | family |
| 22194618 | FP | 2 | l-Cysteine desulfurases | ambiguous name | family |
| 23533635 | FP | 2 | BAG6 complex | ambiguous name | protein |
| 23533635 | FP | 11 | BAG6 | ambiguous name | protein |
| 27751725 | FP | 4 | DDK | ambiguous name | protein |
| 10189366_50 | FP | 1 | kinesin | ambiguous name | family |
| 19415799_9 | FP | 7 | MinD | ambiguous name | protein |
| 19837034_29 | FP | 2 | replisome | ambiguous name | general term |
| 23685356_31 | FP | 1 | POT1 | ambiguous name | protein |
| 23840694_26 | FP | 2 | SNARE | ambiguous name | family |
| 15492226 | FP | 1 | TAK1 kinase complex | annotation error | |

| PubMed ID | FP/ FN | error count | entity | error type | comment |
|---|---|---|---|---|---|
| 21807881 | FP | 1 | COG complex | annotation error | |
| 21807881 | FP | 1 | conserved oligomeric Golgi | annotation error | |
| 21807881 | FP | 4 | COG | annotation error | |
| 25107275 | FP | 1 | clathrin | annotation error | |
| 21700221_12 | FP | 1 | APC/C | annotation error | |
| 11809749 | FP | 1 | polyribosomes | dictionary error | obsolete GO term |
| 13678582 | FP | 1 | cyclin B1-Cdk1 complex | dictionary error | name should be blocked |
| 15492226 | FP | 1 | Ubc13-Uev1A | dictionary error | name should be blocked |
| 19228687 | FP | 1 | MSH2/MSH6 | dictionary error | name should be blocked |
| 19228687 | FP | 1 | MSH2/MSH3 | dictionary error | name should be blocked |
| 20129058 | FP | 1 | DnaB-DnaC complex | dictionary error | name should be blocked |
| 21807881 | FP | 1 | Syntaxin 6-Syntaxin16-Vti1a | dictionary error | name should be blocked |
| 22033972 | FP | 2 | Swi5-Sfr1 | dictionary error | name should be blocked |
| 22033972 | FP | 6 | Swi5-Sfr1 complex | dictionary error | name should be blocked |
| 9857197 | FP | 2 | AP-1 | part-of longer entity | |
| 11521196 | FP | 1 | prefoldin | part-of longer entity | |
| 16912035 | FP | 1 | low density lipoprotein | part-of longer entity | |
| 16912035 | FP | 2 | LDL | part-of longer entity | |
| 17015689 | FP | 1 | IKK | part-of longer entity | |
| 17015689 | FP | 2 | IkappaB kinase | part-of longer entity | |
| 17449811 | FP | 1 | CALCINEURIN | part-of longer entity | |
| 17532339 | FP | 1 | troponin | part-of longer entity | |
| 17532339 | FP | 2 | pyruvate dehydrogenase | part-of longer entity | |

Supplementary Table 6: Error analysis results for transformer-based tagger run on the CoNECo test set.

| PubMed ID | FP/ FN | error count | entity | error type | comment |
|---|---|---|---|---|---|
| 27350684 | FN | 2 | 3'-end processing | ambiguous name | mislabeling; potentially mistaken for biological process |
| 17615152 | FN | 1 | activator protein-1 | ambiguous name | mislabeling; potentially mistaken for family/protein |
| 9367446 | FN | 1 | alpha(v)beta1 | ambiguous name | mislabeling; potentially mistaken for family/protein |
| 19503082 | FN | 4 | AMPAR | ambiguous name | mislabeling; potentially mistaken for family/protein |
| 19637179 | FN | 1 | COX | ambiguous name | mislabeling; potentially mistaken for family/protein |
| 15223 | FN | 1 | cytochrome c oxidase | ambiguous name | mislabeling; potentially mistaken for family/protein |
| 26052893_5 | FN | 1 | ECF transporter | ambiguous name | mislabeling; potentially mistaken for family/protein |
| 26052893_5 | FN | 1 | ECF transporters | ambiguous name | mislabeling; potentially mistaken for family/protein |
| 16024914 | FN | 1 | HDL | ambiguous name | mislabeling; potentially mistaken for family/protein |
| 3453108 | FN | 1 | high density lipoprotein | ambiguous name | mislabeling; potentially mistaken for family/protein |
| 16912035 | FN | 1 | LDL | ambiguous name | mislabeling; potentially mistaken for family/protein |
| 8232104 | FN | 1 | low density lipoprotein | ambiguous name | mislabeling; potentially mistaken for family/protein |
| 24498436 | FN | 3 | Pol V | ambiguous name | mislabeling; potentially mistaken for family/protein |
| 21347277_5 | FN | 3 | RFC | ambiguous name | mislabeling; potentially mistaken for family/protein |
| 18838202 | FN | 9 | TRAIL | ambiguous name | mislabeling; potentially mistaken for family/protein; the long form of the name is weird (tumor necrosis factor-related apoptosis-inducing ligand) and could throw the model off |
| 12815069 | FN | 2 | TRAIL | ambiguous name | mislabeling; potentially mistaken for family/protein |
| 18838202 | FN | 1 | tumor necrosis factor (TNF)-related | ambiguous name | mislabeling; potentially mistaken for family/protein |

| PubMed ID | FP/ FN | error count | entity | error type | comment |
|---|---|---|---|---|---|
| | | | apoptosis-inducing ligand | | |
| 15449939 | FN | 1 | type I human interferon receptor | ambiguous name | mislabeling; potentially mistaken for family/protein |
| 24498436 | FN | 2 | DDR | unidentified model error | |
| 9030599 | FN | 1 | interleukin-5 receptor | unidentified model error | |
| 9030599 | FN | 1 | interleukin-6 receptor | unidentified model error | |
| 17948059 | FN | 7 | Paf1 | unidentified model error | |
| 30802237_18 | FN | 1 | RENT | unidentified model error | |
| 19017259 | FN | 1 | tissue factor-activated FVII | unidentified model error | |
| 1939122 | FP | 1 | 55 S initiation complexes | ambiguous name | ribosome is not a complex according to GO, but consists of complexes |
| 1939122 | FP | 1 | animal mitochondrial ribosomes | ambiguous name | ribosome is not a complex according to GO, but consists of complexes |
| 29666278 | FP | 1 | APC/CCDC20 | ambiguous name | protein |
| 8521815 | FP | 14 | APO-1 | ambiguous name | I think because it’s mentioned as oligomerized at some point |
| 12805216 | FP | 6 | BCR | ambiguous name | protein |
| 8932298 | FP | 1 | casein kinase II | ambiguous name | family |
| 29666278 | FP | 1 | CRL4CDT2 | ambiguous name | protein |
| 17532339 | FP | 1 | dihydrolipoyl acetyltransferase | ambiguous name | family |
| 24768535 | FP | 1 | DNA polymerases | ambiguous name | family |
| 17532339 | FP | 1 | E2 | ambiguous name | family |
| 17572495 | FP | 1 | enhancersome | ambiguous name | general term |
| 9857197 | FP | 1 | export complexes | ambiguous name | general term |

| PubMed ID | FP/ FN | error count | entity | error type | comment |
|---|---|---|---|---|---|
| 22194618 | FP | 1 | formate dehydrogenases | ambiguous name | family |
| 16717095 | FP | 2 | G protein betagamma | ambiguous name | group of complexes |
| 16717095 | FP | 1 | Gbeta | ambiguous name | protein |
| 16717095 | FP | 6 | Gbetagamma | ambiguous name | group of complexes |
| 7524088 | FP | 1 | Gi/Go | ambiguous name | protein |
| 12036871 | FP | 1 | glycoprotein (GP) Ib alpha | ambiguous name | protein |
| 12036871 | FP | 1 | glycoprotein Ib alpha | ambiguous name | protein |
| 12036871 | FP | 2 | GPIb alpha | ambiguous name | protein |
| 18036997 | FP | 1 | immunoglobulin | ambiguous name | family |
| 8407894 | FP | 1 | initiation transcription factor IIIB | ambiguous name | protein |
| 20427570_13 | FP | 1 | malate dehydrogenase | ambiguous name | family |
| 1939122 | FP | 2 | mitochondrial | ambiguous name | discontinuous entity |
| 9362506_22 | FP | 3 | NSF | ambiguous name | protein |
| 24768535 | FP | 10 | PCNA | ambiguous name | protein |
| 25690888 | FP | 4 | PCNA | ambiguous name | protein |
| 25602519 | FP | 3 | PCNA | ambiguous name | protein |
| 20463880 | FP | 1 | PCNA | ambiguous name | protein |
| 16717095 | FP | 1 | phosducin | ambiguous name | group of complexes |
| 17707230 | FP | 1 | polymerases | ambiguous name | family |
| 20129058 | FP | 3 | primase | ambiguous name | family |

| PubMed ID | FP/ FN | error count | entity | error type | comment |
|---|---|---|---|---|---|
| 19837034_29 | FP | 2 | replisome | ambiguous name | general term |
| 15117945 | FP | 1 | ribosome | ambiguous name | ribosome is not a complex according to GO, but consists of complexes |
| 1939122 | FP | 3 | ribosomes | ambiguous name | ribosome is not a complex according to GO, but consists of complexes |
| 29666278 | FP | 1 | SCFSKP2 | ambiguous name | protein |
| 21807881 | FP | 1 | SNARE | ambiguous name | family |
| 8407894 | FP | 1 | transcription factor IIIB | ambiguous name | protein |
| 10866691 | FP | 1 | type I casein | ambiguous name | family |
| 10866691 | FP | 1 | type I casein kinases | ambiguous name | family |
| 25938661_25 | FP | 1 | 3BMP | ambiguous name | PDB ID |
| 25938661_25 | FP | 1 | 4BQ6 | ambiguous name | PDB ID |
| 25380047_22 | FP | 2 | AP-3 | annotation error | |
| 21700221_12 | FP | 1 | APC/C | annotation error | |
| 12805216 | FP | 1 | B-cell antigen receptor | annotation error | |
| 25602519 | FP | 5 | CAF-I | annotation error | |
| 25602519 | FP | 2 | Chromatin Assembly Factor I | annotation error | |
| 25107275 | FP | 1 | clathrin | annotation error | |
| 21807881 | FP | 5 | COG | annotation error | |
| 21807881 | FP | 1 | conserved oligomeric Golgi | annotation error | |
| 30802237 | FP | 5 | Pol I | annotation error | |
| 9115214 | FP | 3 | T cell antigen receptor | annotation error | |

| PubMed ID | FP/ FN | error count | entity | error type | comment |
|---|---|---|---|---|---|
| 9857197 | FP | 2 | AP-1 | part-of longer entity | |
| 17015689 | FP | 1 | IkappaB kinase | part-of longer entity | |
| 17015689 | FP | 1 | IKK | part-of longer entity | |
